# Resistance to pirimiphos-methyl in West African *Anopheles* is spreading via duplication and introgression of the *Ace1* locus

**DOI:** 10.1101/2020.05.18.102343

**Authors:** Xavier Grau-Bové, Eric Lucas, Dimitra Pipini, Emily Rippon, Arjèn van’t Hof, Edi Constant, Samuel Dadzie, Alexander Egyir-Yawson, John Essandoh, Joseph Chabi, Luc Djogbénou, Nicholas J. Harding, Alistair Miles, Dominic Kwiatkowski, Martin J. Donnelly, David Weetman, The *Anopheles gambiae* 1000 Genomes Consortium

## Abstract

Vector population control using insecticides is a key element of current strategies to prevent malaria transmission in Africa. The introduction of effective insecticides, such as the organophosphate pirimiphos-methyl, is essential to overcome the recurrent emergence of resistance driven by the highly diverse *Anopheles* genomes. Here, we use a population genomic approach to investigate the basis of pirimiphos-methyl resistance in the major malaria vectors *Anopheles gambiae* and *A. coluzzii*. A combination of copy number variation and a single non-synonymous substitution in the acetylcholinesterase gene, *Ace1*, provides the key resistance diagnostic in an *A. coluzzii* population from Côte d’Ivoire that we used for sequence-based association mapping, with replication in other West African populations. The *Ace1* and substitution and duplications occur on a unique resistance haplotype that evolved in *A. gambiae* and introgressed into *A. coluzzii*, and is now common in West Africa probably due to cross-resistance with previously used insecticides. Our findings highlight the phenotypic value of this complex resistance haplotype and clarify its evolutionary history, providing tools to understand the current and future effectiveness of pirimiphos-methyl based interventions.

## Introduction

Pirimiphos-methyl is an organophosphate insecticide that is widely used in control interventions against populations of the malaria vector *Anopheles*, especially in Africa (Oxborough 2016; Dengela et al. 2018). Since 2013, the World Health Organization (WHO) has recommended the use of pirimiphos-methyl for indoor residual spraying (IRS) interventions, the major anti-vector strategy in malaria control after treated bednet distribution (Oxborough 2016; World Health Organization 2013). Strategic approaches to vector control often rely on the use of multiple insecticides to avoid or overcome the recurrent emergence of resistance in natural populations (World Health Organization 2018a). In that regard, various insecticide classes have been used in IRS, with pyrethroids—which target the voltage-gated sodium channel—being the dominant choice until recently (Sherrard-Smith et al. 2018; van den Berg et al. 2012). However, the increase in pyrethroid resistance in *Anopheles* populations (Ranson et al. 2011) has led to a progressive replacement with acetylcholinesterase-targeting insecticide classes, first the carbamate bendiocarb, and, latterly, the organophosphate pirimiphos-methyl (Oxborough 2016). Pirimiphos-methyl is the active ingredient in the most widely used insecticide for IRS in Africa, the spray formulation Actellic, which is highly effective and has strong residual performance (Dengela et al. 2018; Sherrard-Smith et al. 2018; Wagman et al. 2018; Abong’o et al. 2020). However, resistance has recently been reported in several populations of African *Anopheles* s.l. (Kisinza et al. 2017; Chukwuekezie et al. 2020), and though control failures have yet to be reported, it represents a clear threat to the efficacy of IRS strategies.

Mechanisms of resistance to pirimiphos-methyl are poorly understood, but organophosphates, as well as carbamates, all block the action of the acetylcholinesterase enzyme (ACE1, encoded by the *Ace1* gene in mosquitoes) via competitive binding to its active site (Oakeshott et al. 2005). Studies in culicine and anopheline mosquitoes have found that non-synonymous mutations in *Ace1* can result in resistance to carbamates and organophosphates other than pirimiphos-methyl (Weill et al. 2003, 2004; Feyereisen et al. 2015). The most common mutation in *Anopheles* is a glycine to serine mutation in codon 280 (*G280S*, also known as *G119S* after the codon numbering of a partial crystal structure from the electric ray *Torpedo californica* (Weill et al. 2003; Greenblatt et al. 2004; Feyereisen et al. 2015)), which is located near the active site gorge of ACE1 (Cheung et al. 2018) and decreases its sensitivity to organophosphates and carbamates. This mutation results in reduced sensitivity to the acetylcholine neurotransmitter (Bourguet et al. 1997), thus carrying potential fitness costs, as demonstrated in *Culex pipiens* (Labbé et al. 2007). The resistance allele *280S* has been regularly found in natural *Anopheles* populations: *A. gambiae s*.*s*. (henceforth, *A. gambiae*) and *A. coluzzii* from West Africa, ranging from Benin to Côte d’Ivoire (Weill et al. 2003; Dabiré et al. 2009; Ahoua Alou et al. 2010; Essandoh et al. 2013; Weetman et al. 2015); as well as in *A. albimanus* (Liebman et al. 2015), *A. arabiensis* (Dabiré et al. 2014), and *A. sinensis* (Feng et al. 2015). In addition, *Ace1* is subject to duplication polymorphisms (copy number variants, or CNVs) that co-segregate with the *280S* allele in both *A. gambiae* and *A. coluzzii*, and enhance resistance to carbamates and organophosphates such as fenitrothion and chlorpyrifos-methyl (Djogbénou et al. 2008; Essandoh et al. 2013; Edi et al. 2014a; Weetman et al. 2015; Assogba et al. 2015, 2016, 2018).

In this study we provide an in-depth investigation of the relationship between *Ace1* mutations and pirimiphos methyl resistance in *A. gambiae* and *A. coluzzii* using whole-genome sequenced samples from the *Anopheles gambiae* 1000 Genomes project (The *Anopheles gambiae* 1000 Genomes Consortium 2019; Miles et al. 2017), and a wider testing of phenotyped specimens from across West Africa. In addition, we perform a first agnostic genome-wide scan for candidate regions contributing to pirimiphos-methyl resistance in a population of *A. coluzzii* from Côte d’Ivoire. Finally, we study the contemporary evolution of *Ace1* to answer unresolved questions on the selective pressures and the pattern of inter-specific introgression associated with the spread of this resistance mechanism. Earlier studies provided support for the idea that a common resistance haplotype under positive selection might have introgressed between species (Djogbénou et al. 2008; Essandoh et al. 2013; Weetman et al. 2015), but they focused on a partial region of the *Ace1* gene and did not address the relationship between introgression and the duplication, which extends beyond the gene (Assogba et al. 2016; Lucas et al. 2019). Here we leverage population genomic resources from the *Anopheles gambiae* 1000 Genomes to overcome these limitations. Overall, our results demonstrate a widespread and dominant role for *Ace1* mutations in pirimiphos-methyl resistance, provide critical insights into resistance diagnosis, and demonstrate that *280S* alleles and *Ace1* duplications co-occur on a single, swept, resistance haplotype that originated in West African *A. gambiae* and later introgressed into *A. coluzzii*.

## Results

### Conservation and distribution of *Ace1* resistance mutations in *Anopheles*

We examined the frequency and distribution of the two *Ace1* mutations that have been associated with organophosphate and carbamate resistance in *A. gambiae* and *A. coluzzii*: the *G280S* non-synonymous single nucleotide polymorphism (SNP), and copy number variation (CNV) polymorphisms of *Ace1* and the surrounding genomic region. *G280S* is sometimes known as *G119S* (Feyereisen et al. 2015) based on its position in the truncated crystal structure of its homolog in the electric ray *Torpedo californica*, where ACE1 protein structure was first elucidated (Greenblatt et al. 2004). Due to a culicine-specific N-terminal insertion in ACE1, the exact position of this conserved codon differs among animal orthologs (Supplementary Material SM1). We provide a list of homologous codon positions for *Ace1* orthologs from selected animal species, including common insect vectors (Supplementary Material SM2). Henceforth, we will use *A. gambiae* s.l.-based codon coordinates and refer to this SNP as *G280S* (wild-type allele: *280G*; resistance allele: *280S*; gene accession number: AGAP001356-RA in AgamP4.12).

In the *Anopheles* 1000 Genomes cohort (Phase 2, *n* = 1142 genomes; Figure 1) (The *Anopheles gambiae* 1000 Genomes Consortium 2019), the *280S* resistance allele is present across West African populations of *A. coluzzii* (Côte d’Ivoire, Burkina Faso, Ghana) and *A. gambiae* (Burkina Faso, Ghana, Guinea), with the highest frequencies being attained in Ghanaian *A. gambiae* (67%) and Ivorian *A. coluzzii* (44%; Figure 1A), which is consistent with previous results (Djogbénou et al. 2008; Dabiré et al. 2009; Ahoua Alou et al. 2010; Essandoh et al. 2013). *Ace1* CNVs have a similar distribution to the SNP in West African populations, as they overwhelmingly overlap with specimens also carrying *280S* alleles (Figure 1B).

**Figure 1.**
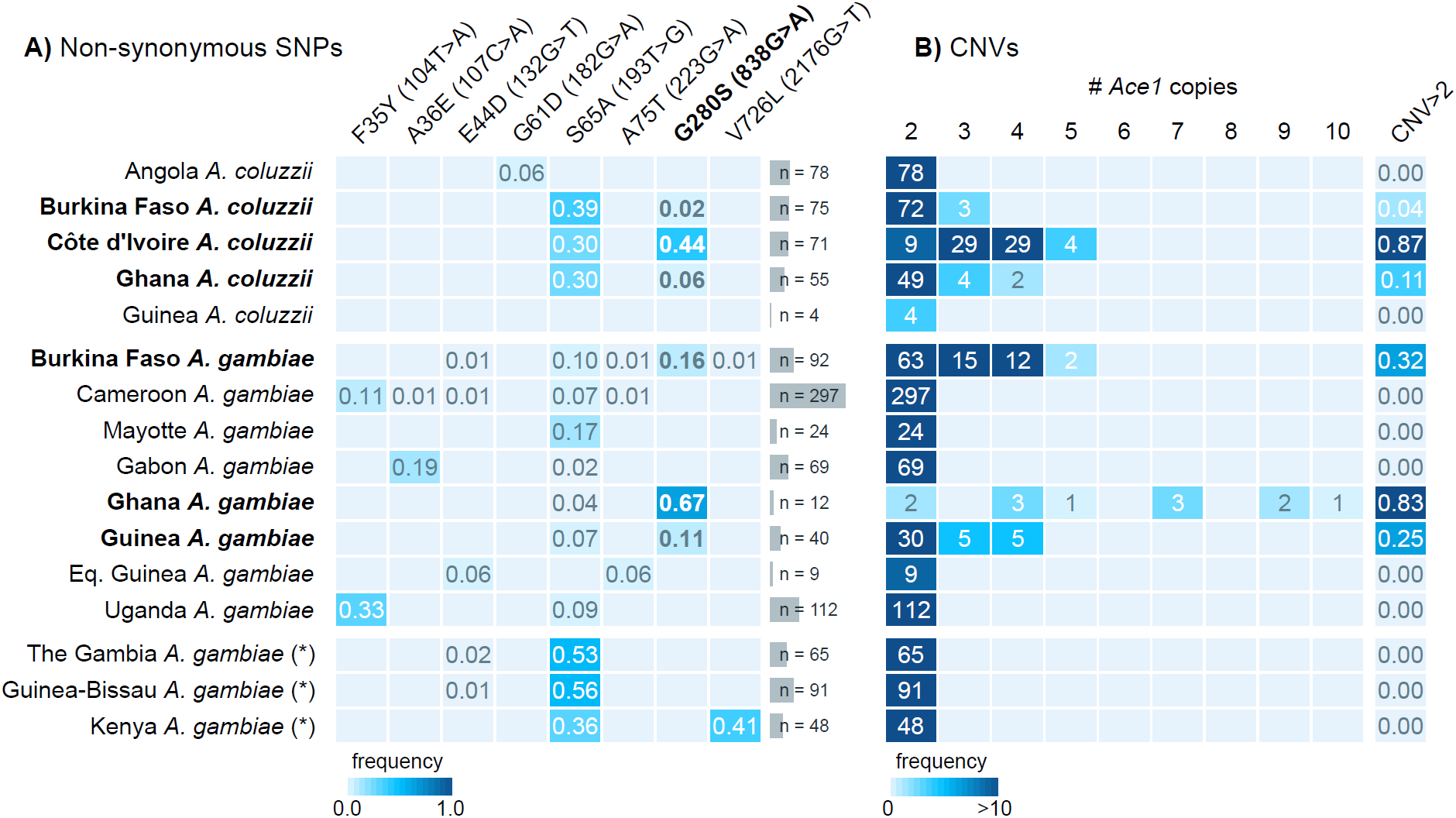
*Ace1* mutations in African populations. **A)** Frequency of non-synonymous SNPs in the *Ace1* gene in African *A. gambiae* and *A. coluzzii* populations from *Anopheles gambiae* 1000 Genomes, Phase 2. For each SNP, we indicate peptide- and transcript-level coordinates and substitutions. **B)** *Ace1* CNVs across African populations, including the frequency of specimens with >2 copies in each population. A diploid genome without duplications would have 2 copies. Populations where *G280S* and duplications are present are highlighted in bold text. Note: populations denoted with an asterisk (The Gambia, Guinea-Bissau and Kenya) have high frequency of hybridisation and/or unclear species identification.

The combination of CNVs and *280S* alleles results in multiple genotypes being observed in the *Ace1* locus, defined by a variable number of *280S* copies. We used the fraction of sequencing reads supporting the *wt* and *280S* alleles to estimate the number of *280S* alleles in each sample (Figure 2A and 2B), which revealed that most specimens with duplications carry both *wt* and *280S* alleles at various frequencies (*n* = 113, orange box in Figure 2A), whereas the vast majority of non-duplicated specimens only have *wt* alleles (*n* = 1021, purple box in Figure 2A). In addition, we identify six high-copy, *280S* homozygous specimens, present only in the Ghanaian *A. gambiae* population in this dataset (Figure 1B).

**Figure 2.**
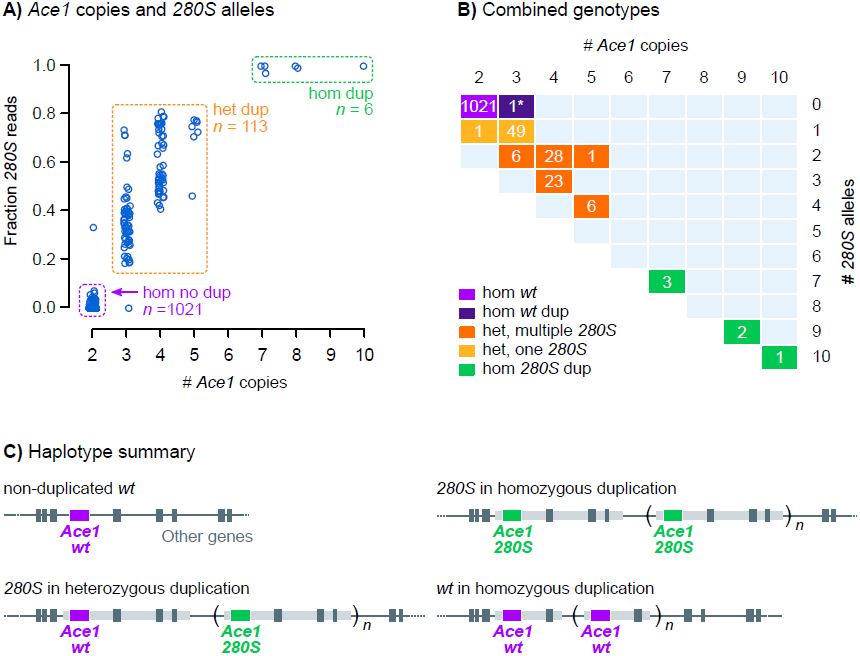
Combinations of *Ace1 G280S* and CNV genotypes. **A)** Fraction of reads supporting *280S* alleles and number of *Ace1* copies (1000 Genomes dataset, *n* = 1142). Boxes highlight groups of haplotypes: non-duplicated *wt* homozygotes (purple), duplicated heterozygotes (orange) and duplicated *280S* homozygotes (green). **B)** Breakdown of of possible genotypes at the *Ace1* locus according to *Ace1* and *280S* copy number. The asterisk (*) denotes a *wt-*homozygous specimen from Guinea that carries an independently evolved *Ace1* duplication. **C)** Summary of haplotypes observed in the *Ace1* locus.

Virtually all *Ace1* CNVs identified in the 1000 Genomes cohort share the same duplication breakpoints (Lucas et al. 2019), spanning a region *ca*. 200 kbp that includes a total of 11 genes (Supplementary Material SM5). The only exception is a single *A. gambiae* specimen from Guinea that carries a unique *wt*-homozygous duplication (asterisk in Figure 2B). This Guinea-specific CNV is shorter than the *Ace1* duplication found in all other samples and has different breakpoints (*ca*. 70 kbp, including *Ace1* and one other gene; Supplementary Material SM4), implying an independent origin. Henceforth, all mentions of *Ace1* CNVs will refer to the major duplication. The absence of specimens carrying the major duplication and lacking *280S* alleles strongly implies that this CNV contains both *280S* and *wt* alleles on the same chromosome, and thus results in permanent heterozygosity. Figure 2C summarises the four model haplotypes that can result in the genotypes observed in the 1000 Genomes dataset: (i) non-duplicated *wt*; (ii) heterozygous and (iii) *280S-* homozygous major duplications; and (iv) the *wt* minor duplication from Guinea.

In addition to *G280S*, we found seven non-synonymous SNPs in *Ace1* with at least 1% frequency in at least one population (Figure 1A). None were in linkage disequilibrium with *G280S* (Supplementary Material SM3), nor have any of them been previously associated with resistance (Feyereisen et al. 2015). The most salient of these additional mutations was a serine-to-alanine SNP in codon 65 (*S65A*), with frequencies of 30-50% in multiple West African populations. Codon 65 maps to the anopheline-specific N-terminal insertion of ACE1 proteins (Supplementary Material SM1), which lacks predicted secondary structure and is far away from the active-site gorge of the enzyme (Cheung et al. 2018). This position is not conserved across ACE1 orthologs, being variously encoded as alanine, serine or threonine in different *Anopheles* species (Supplementary Material SM1).

### Pirimiphos-methyl resistance in Ivorian *A. coluzzii* is linked to *Ace1* duplications and multiple *280S* alleles

Next, we examined the association between *Ace1* mutations and resistance to pirimiphos-methyl (Figure 3). We used 71 *A. coluzzii* female mosquitoes from Côte d’Ivoire, collected from rice fields in Tiassalé in 2012, that had been tested for resistance prior to genome sequencing. The *G280S* SNP and CNV co-occur at high frequencies in this population, with 87.3% of specimens being duplicated heterozygotes with a one or more *280S* alleles (Figure 3A). Specimens with at least one *280S* allele had a survival rate of 50%, as opposed to 0% in the *wt* homozygotes (Figure 3B), suggesting that *280S* is linked to pirimiphos-methyl resistance in this population (*p =* 3.7 × 10^−3^ in Fisher’s exact test, Woolf-corrected odds ratio = 19.0 [95% confidence interval = 1.1 – 340.6]) but it does not fully explain the resistance phenotype. The only other non-synonymous SNP present in Ivorian *A. coluzzii, S65A*, was not associated with pirimiphos-methyl resistance (*p =* 0.605 Fisher’s exact test; Supplementary Material SM6A, SM6B).

**Figure 3.**
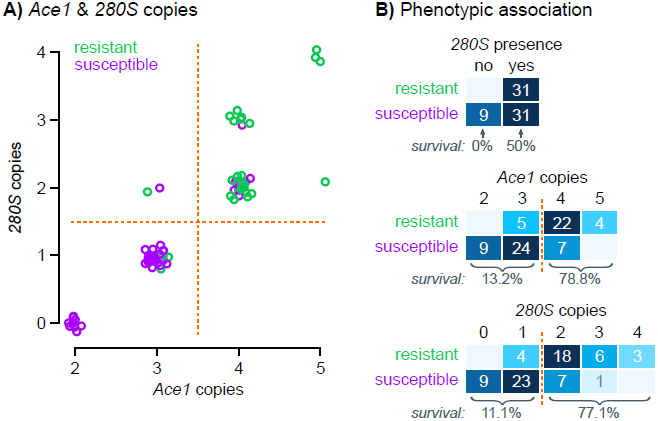
Genotype-phenotype association in Ivorian *A. coluzzii*. **A)** Number of *Ace1* copies compared to the estimated number of *280S* copies in Ivorian *A. coluzzii* (n=71), color-coded according to resistance phenotypes. Random jitter has been added for clarity. **B)** Cross-tabulation of pirimiphos-methyl resistance and three *Ace1* mutations in Ivorian *A. coluzzii*: *280S* allele presence, number of *Ace1* copies, and number of *280S* alleles. Orange lines denote *ad hoc* groups of genotypes where we identify changes in survival rates (included at the bottom of each table). Source data available in Supplementary Material SM6.

We found a strong bias towards higher *Ace1* copy numbers in resistant specimens, all of which had at least 3 copies of the *Ace1* locus and most of them had 4 or more (Figure 3B; odds ratio from a binomial generalised linear model [GLM] = 16.9 [5.1 – 56.6]; *p* = 6.4 × 10^−10^ in a *χ*^*2*^ test compared to a null model). Concordantly, the estimated number of *280S* alleles was also associated with resistance (GLM odds ratio = 10.7 [3.6 – 31.7]; *p* = 8.9 × 10^−10^ in *χ*^*2*^ test): survival rates increased with higher number of *280S* copies (Figure 3B), with a threshold apparent between individuals with zero or one copies of *280S* (11.1% survival) and those with two or more (77% survival). Crucially, within the group of duplicated specimens with one *280S* and two *wt* alleles (*n* = 36), *280S* provided no survival benefit compared to non-duplicated *wt* specimens (Fisher’s exact test *p =* 0.553; Woolf-corrected odds ratio = 3.6 [0.2 – 74.3]).

We also examined the combined effects of all the mutations detected in the *Ace1* locus (*280S* and *S65A* presence, CNVs and number of *280S* alleles) in Ivorian *A. coluzzii*. We used a binomial GLM and a step-wise procedure to remove non-informative variants according to the Bayesian Information Criterion (see Methods), finding that the number of *Ace1* copies was the minimal sufficient marker to predict pirimiphos-methyl resistance in this population (odds ratio = 16.9 [5.1 – 56.9]; *p* = 6.43 × 10^−10^ in a *χ*^*2*^ test; details in Supplementary Material SM6B).

### Pirimiphos-methyl resistance diagnostics are repeatable across *A. gambiae* and *A. coluzzii* populations

We investigated the extent to which the role of *Ace1* and *280S* copy numbers in pirimiphos-methyl resistance could be generalised to other populations beyond the *A. coluzzii* from Tiassalé. Specifically, we surveyed *Ace1* mutations and phenotypic resistance in eight West African populations of *A. gambiae* and *A. coluzzii* from additional locations in Côte d’Ivoire, Togo, and Ghana (Table 1 and Table 2; list of specimens in Supplementary Material SM6C and SM6D).

**Table 1.** Association of resistance genotypes with pirimiphos-methyl resistance in nine West African populations of *A. coluzzii* and *A. gambiae*. For each population, we report number of sampled specimens (*n*), the number of *wt* homozygous (*wt*), heterozygous (*het*) and *280S* homozygous specimens (*hom*), survival frequency among mutated groups, and the odds ratios of survival in heterozygous and *280S* homozygous specimens (if available) in a binomial generalised linear model, and the *p-*value of this model in a *χ*^*2*^ ANOVA comparison with a null model. Odds ratios (OR) are reported with 95% confidence intervals. Dagger signs (†) denote survival rates in genotypes with less than 5 specimens. Samples listed in Supplementary Material SM6C; statistical analysis in Supplementary Material SM6D. Country abbreviations: CI, Côte d’Ivoire; GH, Ghana; TG, Togo.

| Population | *n* | Frequencies<br>*wt* / *het* / *hom* | % survival<br>*het* | OR<br>*het* | % survival<br>*hom* | OR<br>*hom* | *p* |
| --- | --- | --- | --- | --- | --- | --- | --- |
| *coluzzii* Aboisso (CI) | 55 | 42 / 13 / 0 | 46.1% | 35.1<br>(3.6 – 338.0) | - | - | $1.3 \times 10^{-4}$ |
| *coluzzii* Korle-Bu (GH) | 214 | 25 / 165 / 24 | 49.6% | $4.2 \times 10^7$<br>(0 – Inf) | 95.8% | $9.8 \times 10^8$<br>(0 – Inf) | $1.2 \times 10^{-13}$ |
| *coluzzii* Weija (GH) | 131 | 60 / 70 / 1 | 54.1% | 7.3<br>(3.0 – 17.6) | 100% (†) | $3.7 \times 10^7$<br>(0 – Inf) | $2.4 \times 10^{-6}$ |
| *gambiae* Aboisso (CI) | 82 | 3 / 45 / 34 | 77.8% | 7.0<br>(0.6 – 85.4) | 97% | 66.0<br>(2.9 – 1491.2) | $3.7 \times 10^{-3}$ |
| *gambiae* Madina (GH) | 172 | 19 / 112 / 41 | 58.9% | 12.2<br>(2.7 – 55.4) | 100% | $9.8 \times 10^8$<br>(0 – Inf) | $4.4 \times 10^{-14}$ |
| *gambiae* Obuasi (GH) | 140 | 0 / 28 / 112 | 17.8% | - | 58.9% | 6.6<br>(2.3 – 18.6) | $6.0 \times 10^{-5}$ |
| *gambiae* Weija (GH) | 13 | 7 / 4 / 2 | 75% (†) | $7.3 \times 10^{17}$<br>(0 – Inf) | 100% (†) | $2.6 \times 10^9$<br>(0 – Inf) | $1.6 \times 10^{-3}$ |
| *gambiae* Baguida (TG) | 102 | 0 / 14 / 88 | 57.1% | - | 56.8% | 0.9<br>(0.3 – 3.1) | 0.982 |

**Table 2.** Association of *Ace1* duplications with pirimiphos-methyl resistance in heterozygous specimens from seven West African populations of *A. coluzzii* and *A. gambiae*. For each population, we report number of sampled specimens (*n*), and the results of three binomial generalised linear models (GLM): (i) using the number of *Ace1* copies as the predictor variable, (ii) using the *280S*-to-*280G* allelic ratio, and (iii) a minimal model obtained with step-wise reduction, according to the Bayesian Information Criterion (BIC). For each model, we report the *p*-value from a comparison with a null model (ANOVA *χ*^*2*^ test) and odds ratios (OR) with 95% confidence intervals for the variables included in the model. Sample phenotypes and genotypes listed in Supplementary Material SM6E; statistical analysis in Supplementary Material SM6F. Country abbreviations: CI, Côte d’Ivoire; GH, Ghana; TG, Togo.

| Population | *n* | GLM *Ace1* copies<br>(single variable) |  | GLM 280S/280G allelic ratio<br>(single variable) |  | Minimal GLM (BIC) |  |  |
| --- | --- | --- | --- | --- | --- | --- | --- | --- |
|  |  | OR | *p* | OR | *p* | OR<br>*Ace1* copies | OR<br>allelic ratio | *p* |
| *coluzzii* Aboisso (CI) | 11 | 1.3<br>(0.7 – 2.3) | 0.361 | 52.9<br>(0.01 –<br>2.2 × 10<sup>5</sup>) | 1.3 × 10<sup>-3</sup> | - | 52.9<br>(0.01 –<br>2.2 × 10<sup>5</sup>) | 1.3 × 10<sup>-3</sup> |
| *coluzzii* Korle-Bu (GH) | 46 | 1.8<br>(1.2 – 2.6) | 1.0 × 10<sup>-4</sup> | 2.8<br>(1.6 – 5.1) | 4.1 × 10<sup>-7</sup> | 1.5<br>(0.9 – 2.3) | 2.3<br>(1.3 – 4.0) | 3.0 × 10<sup>-7</sup> |
| *coluzzii* Weija (GH) | 21 | 0.5<br>(0.3 – 1.0) | 0.013 | 3.4<br>(0.4 – 28.6) | 0.021 | 0.4<br>(0.1 – 1.1) | 50<br>(0.3 – 9505.0) | 2.9 × 10<sup>-3</sup> |
| *gambiae* Aboisso (CI) | 31 | 1.0<br>(0.6 – 1.5) | 0.933 | 2.6<br>(0.3 – 24.9) | 0.382 | - | - | - |
| *gambiae* Madina (GH) | 19 | 5.2<br>(1.1 – 23.9) | 7.7 × 10<sup>-3</sup> | 6.5 × 10<sup>85</sup><br>(0 – Inf) | 2.1 × 10<sup>-6</sup> | 8.7 × 10<sup>-20</sup><br>(0 – Inf) | 5.5 × 10<sup>45</sup><br>(0 – Inf) | 2.6 × 10<sup>-9</sup> |
| *gambiae* Obuasi (GH) | 26 | 1.5<br>(0.9 – 2.3) | 0.038 | 3.1<br>(0.85 – 10.8) | 0.038 | 1.5<br>(0.9 – 2.3) | - | 0.038 |
| *gambiae* Weija (GH) | 0 | - | - | - | - | - | - | - |
| *gambiae* Baguida (TG) | 13 | 1.4<br>(0.7 – 2.6) | 0.350 | 2.9<br>(0.1 – 69.4) | 0.498 | - | - | - |

Pirimiphos-methyl resistance was associated with *G280S* mutations in seven out of eight populations (*p <* 0.01 in a *χ*^*2*^ test comparing GLM of each genotype to a null model; Table 1). However, whilst *280S* homozygotes were strongly associated with resistance in populations where they were present in significant numbers (e.g. 95.8% survival rate in *A. coluzzii* from Korle-Bu), heterozygotes often exhibited lower, intermediate survival rates (e.g. 49.6% in *A. coluzzii* from Korle-Bu; six out of eight populations had survival rates <60%; Table 1) similar to that of the *A. coluzzii* population from Tiassalé (Côte d’Ivoire) mentioned above (Figure 3A).

Following our findings for the Tiassalé population (Figure 3), we further investigated the phenotypic variation in duplicated heterozygotes using two CNV-related variables: (i) the ratio of *280S* to *280G* alleles (measured as the ratio of FAM-to-HEX dye signal in a TaqMan qPCR assay) (Bass et al. 2010); and (ii) the estimated number of *Ace1* copies relative to a non-duplicated and *wt* Kisumu *A. gambiae* colony (Edi et al. 2014a) (see Methods).

Pirimiphos-methyl resistance was associated with CNV-related measures in heterozygotes from *A. gambiae* and *A. coluzzii* populations, according to generalised linear models that included *280S* allelic ratios and/or *Ace1* copy number as potential informative markers (Table 2). In *A. coluzzii* from Korle-Bu and Weija, as well as *A. gambiae* from Madina, both the *280S* allelic ratio and the *Ace1* copy number were included in the minimal model (at *p <* 0.01 in a *χ*^*2*^ comparison with a null model); whereas in *A. coluzzii* from Aboisso the allelic ratio was the only variable associated with resistance (*p* = 1.3 × 10^−3^ in a *χ*^*2*^). In the Obuasi *A. gambiae* heterozygotes, the minimal model only included *Ace1* copy number (at *p* = 0.038) but neither measure was strongly predictive, as was the case in *A. gambiae* from Aboisso and Baguida. Overall, these results indicate that the combination of *280S* allelic ratios and CNVs provide a similar pirimiphos-methyl resistance mechanism in heterozygotes from both species. However, diagnostic capacity appears to vary among populations, suggesting the possible existence of additional resistance mechanisms.

### Genome-wide identification of pirimiphos-methyl resistance variants in Ivorian *A. coluzzii*

We examined the genomes of the 71 *A. coluzzii* specimens from Côte d’Ivoire to identify additional genetic variants—other than *Ace1 G280S* and duplications—that could be linked to pirimiphos-methyl resistance. All Ivorian specimens had been collected in the same location and time (Tiassalé, 2012), before the widespread adoption of pirimiphos-methyl in IRS interventions (Oxborough 2016; World Health Organization 2013). The absence of a long period of pirimiphos-methyl use prior to collection implies that adaptation to this insecticide would presumably be caused by cross-resistance with previously employed insecticides and/or standing variation, rather than novel selective sweeps. In accordance with this expectation of low differentiation, a principal component analysis of genetically unlinked variants (see Methods) did not reveal population stratification between resistant and susceptible mosquitoes (Figure 4), and average Hudson’s *F*_*ST*_ between them was low along all chromosomal arms (*F*_*ST*_ *≈* 0%).

**Figure 4.**
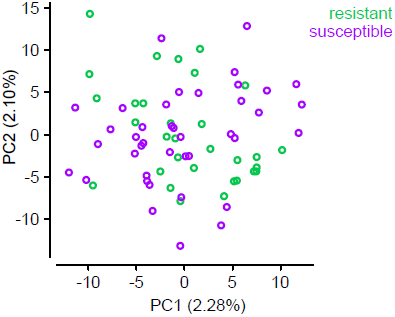
Principal component analysis of Côte d’Ivoire *A. coluzzii*. PCA constructed using genotypes of 791 unlinked variants from chromosome 3.

We thus aimed to identify signals of selection in resistant *A. coluzzii* using the population branch statistic (*PBS*, Figure 5A), which identifies regions with an excess of genetic differentiation in a focal population (here, resistant Ivorian mosquitoes) relative to a basal level of differentiation between a closely related population (susceptible Ivorian mosquitoes) and a more distant control (a population of *A. coluzzii* from Angola; see Methods). *PBS* is a powerful test to detect recent selection acting on incomplete sweeps and standing variation (Yi et al. 2010; Malaspinas et al. 2016), and is therefore well-suited to investigate a scenario—like ours—in which the population of interest has not yet diverged.

**Figure 5.**
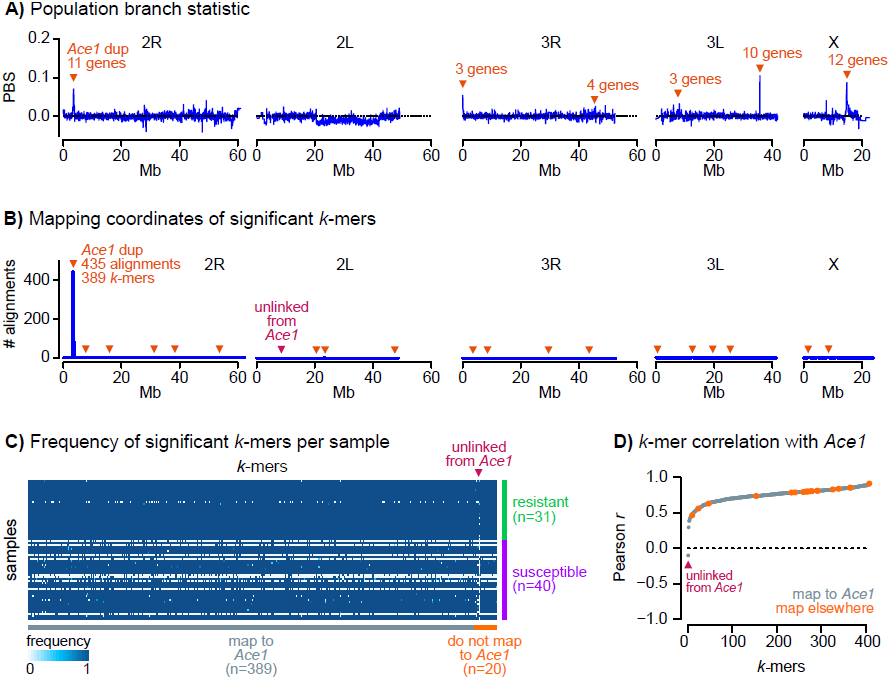
Genome-wide scan of variants associated with pirimiphos-methyl resistance in Ivorian *A. coluzzii*. **A)** Profile of population branching statistics along all chromosomal arms, calculated in consecutive blocks of 1000 segregating variants, using resistant and susceptible Ivorian *A. coluzzii* as populations A and B, and Angolan *A. coluzzii* as outgroup. Orange triangles indicate windows with extreme *PBS* values (*p-*values derived from a standardised distribution of PBS along each chromosomal arm, and *FDR* < 0.001), and the number of genes therein. **B)** Mapping coordinates of *k-*mers significantly associated with pirimiphos-methyl (*n*=439). Most *k*-mers map to the *Ace1* duplication region (*n*=414) or, despite mapping elsewhere in the genome (*n*=24), are correlated with *Ace1* copy number (orange triangles). Only one *k-*mer mapping outside of the *Ace1* duplication is not correlated with *Ace1* copy number (purple triangle). **C)** Normalised frequency of each significant *k*-mer (*n*=439, horizontal axis) in each genome (*n*=71, vertical axis). *k*-mers are sorted according to their mapping location (in *Ace1* or elsewhere), and genomes are sorted according to their phenotype (resistant/susceptible). **D)** Pearson’s correlation coefficients (*r*) between *k-*mer frequency and number of *Ace1* copies in each genome (*n*=439 significant *k-*mers). *k*-mers are coloured according to their mapping location (in *Ace1* or elsewhere) and sorted by the values of *r*.

We estimated *PBS* along genomic windows to identify regions in which resistant Ivorian specimens had an excess of genetic differentiation (see Methods). In total, we identified 43 genes within six regions with excess differentiation in resistant *A. coluzzii* (*PBS* > 0, *FDR* < 0.001; see Methods; Figure 5A). This set of candidates contained eleven mostly uncharacterised genes located downstream of *Ace1* and within the duplicated region, two ionotropic receptors (*GLURIIb* and *GLURIIa*), and several protesases, among others (Supplementary Material SM7). However, genetic differentiation between resistant and susceptible specimens in each of these genes was low (*F*_*ST*_ < 6% in all cases; Supplementary Material SM7), which suggests that they are unlikely to be directly associated with the resistance phenotype. The *Ace1* gene itself was on the verge of the significance threshold (*FDR* = 3.2 × 10^−3^, *PBS* = 0.031) and exhibited low differentiation (*F*_*ST*_ = 1.2%). This apparent contradiction regarding the *Ace1* phenotypic association can be explained by the fact that pirimiphos-methyl resistance is caused by a combination of mutations—a SNP within a heterozygous duplication—that is not captured by the diploid allelic frequencies used in *PBS* calculations (Yi et al. 2010).

To overcome this limitation, we performed a *k-*mer association study comparing the resistant and susceptible Ivorian *A. coluzzii. k*-mer association studies test for differential frequencies of tracts of nucleotides of length *k* in different groups of genomes, and are able to identify genetic variation patterns linked to both SNPs and structural variants, such as CNVs, without requiring a reference annotation (Rahman et al. 2018). We identified *ca*. 767 million different *k*-mers of length *k* = 31 bp across all 71 samples (see Methods). Among these, 9,603 *k*-mers were significantly enriched in resistant specimens (Spearman correlation test, *FDR* < 0.001). The 9,603 significant *k-*mers were assembled into 446 unique sequences composed of more than one *k*-mer (median length = 54 bp), which we then aligned to the *A. gambiae* reference genome. We retained sequences that could be mapped to chromosomes 2, 3 or X (409 out of 446) for further analysis (listed in Supplementary Material SM8). Among these, the vast majority (*n* = 389, 95.1%) aligned to the *Ace1* duplicated region, while the remaining significant sequences aligned in scattered regions across the rest of the genome (*n* = 20, 4.9%; Figure 5B). Yet, 19 out of 20 *k-*mers that did not map to *Ace1* had a very similar frequency profile in resistant and susceptible samples than the 389 *Ace1*-linked *k*-mers, being absent and present in the same genomes (Figure 5C). In fact, the *k*-mer frequencies of these 19 sequences was strongly correlated with *Ace1* copy number (Pearson correlation *r* > 0 and *p* < 1 × 10^−4^; Figure 5D). This suggests that these 19 *k*-mers represent a non-independent signal that also reflects the association of *Ace1* mutations with resistance, and can be parsimoniously explained by non-specific sequence alignments (see Discussion). Only one remaining sequence did not correlate with *Ace1* copy number (Pearson correlation *r* < 0.0, maroon arrow in Figure 5C and 5D), indicating that it may represent another variant involved in resistance. The primary alignment of this sequence mapped to a non-coding region of chromosomal arm 2L (from 2L:8,662,023), with no proximity to any gene of known function. Altogether, these results support the conclusion that *Ace1* mutations are the primary driver of pirimiphos-methyl resistance in this *A. coluzzii* population.

### Selection and introgression of *Ace1* duplications in *A. gambiae* and *A. coluzzii*

A high degree of geographical and phylogenetic overlap is evident between *Ace1* duplications and *280S* alleles across four countries (Côte d’Ivoire, Ghana, Burkina Faso and Guinea) and two different species (*A. coluzzii* and *A. gambiae*; Figure 2A). Previous studies provide two key insights to understand this pattern. First, *Ace1* duplications across West African populations have concordant breakpoints (Assogba et al. 2016; Lucas et al. 2019), which suggests they share a common evolutionary origin despite their multi-species distribution. Second, partial sequencing of *Ace1* has shown that *280S* alleles share highly similar haplotypic backgrounds in both West African *A. gambiae* and *A. coluzzii* (Djogbénou et al. 2008; Essandoh et al. 2013; Weetman et al. 2015), which is suggestive of inter-specific introgression and a selective sweep. The most parsimonious hypothesis linking our results and these observations would posit that (i) the high similarity of *280S*-carrying haplotypes across the *A. gambiae – A. coluzzii* species boundary is shared along the entire duplicated region (*ca*. 200 kbp); and (ii) this similarity is due to a *280S-*linked selective sweep around *Ace1*, followed by duplications and the joint introgression of the duplication-*280S* haplotype between *A. gambiae* and *A. coluzzii*.

To test these hypotheses, we investigated genetic variation in the *Ace1* duplication using networks of haplotype similarity (Figure 6). The analysis of *Ace1* haplotypes is limited by the low density of phased variants in the vicinity of this gene (Supplementary Material SM9), caused by a combination of high sequence conservation and the difficulties in phasing haplotypes within a heterozygous duplication. To circumvent these limitations, we built four different networks around tagging variants that were in strong linkage disequilibrium with *G280S* (Huff and Rogers’ *r* > 0.95) and located within the duplication (located within −26 kbp and +12 kbp from *G280S*; Supplementary Material SM9), and used these to define four sets of *280S*-linked haplotypes. Then, we assumed that concordant observations among all sets of *280S*-linked haplotypes would reflect genetic signatures shared with *Ace1*.

**Figure 6.**
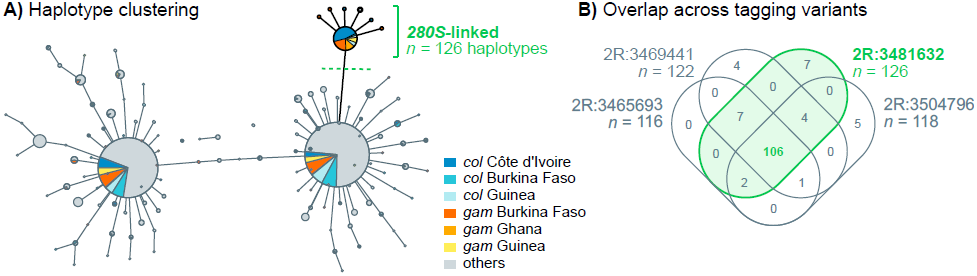
Haplotype clustering network around *Ace1*. **A)** Minimum spanning tree of haplotypes around the *280S-*linked variant 2R:3481632 (± 300 bp, *n* = 104 phased variants). Node size reflects number of haplotypes belonging to each cluster, which are color-coded according to species and geographical origin. Edges link haplotype clusters separated by one substitution. Singleton clusters have been removed from this view (see Supplementary Material SM10). A cluster of identical haplotypes linked to the *280S-*tagging variant is highlighted in green. **B)** Venn diagram representing the overlap of samples belonging to the *280S-*linked haplotype clusters identified around each of the four tagging variants (Supplementary Material SM10).

The minimum spanning tree built from haplotypes located around the first tagging variant (Figure 6A) identified a cluster of 124 identical haplotypes carrying the *280S*-linked allele (Figure 6A, labelled with green text) and two larger haplotype clusters linked to the *wt 280G* allele (Figure 6A). The *280S* cluster contains identical haplotypes from West African populations (Côte d’Ivoire, Burkina Faso, Ghana and Guinea) of both *A. coluzzii* and *A. gambiae*. On the other hand, *wt*-linked clusters have wider geographical distributions (including samples from West, Central and Eastern sub-Saharan Africa). The other three tagging variants show haplotype networks with a similar structure, with clusters of *ca*. 120 *280S*-linked identical haplotypes originating in the same *A. gambiae* and *A. coluzzii* specimens from West Africa (Figure 6B, Supplementary Material SM10). This result indicates that haplotype similarity between *A. gambiae* and *A. coluzzii* extends beyond the *Ace1* gene.

We also investigated possible signals of positive selection around the *Ace1* duplication breakpoints using Garud’s *H* statistics, haplotypic diversity, and extended haplotype homozygosity (Figure 7, Supplementary Material SM11 and SM12). The profile of Garud’s *H*_*12*_ statistic in *280S-*linked haplotypes showed peaks both upstream and downstream of the *Ace1* duplication (Figure 7A), which coincided with a low *H*_*2*_*/H*_*1*_ ratio (Figure 7B) and low haplotypic diversity (Figure 7C), and is thus indicative of a hard selective sweep in this region (Garud et al. 2015; Messer and Petrov 2013). Notably, the region of extended haplotype homozygosity associated with this sweep was still apparent upstream and downstream of the duplication breakpoints, i.e., far away from the focal tagging variant used to define *280S*-linked haplotypes (Figure 7D).

**Figure 7.**
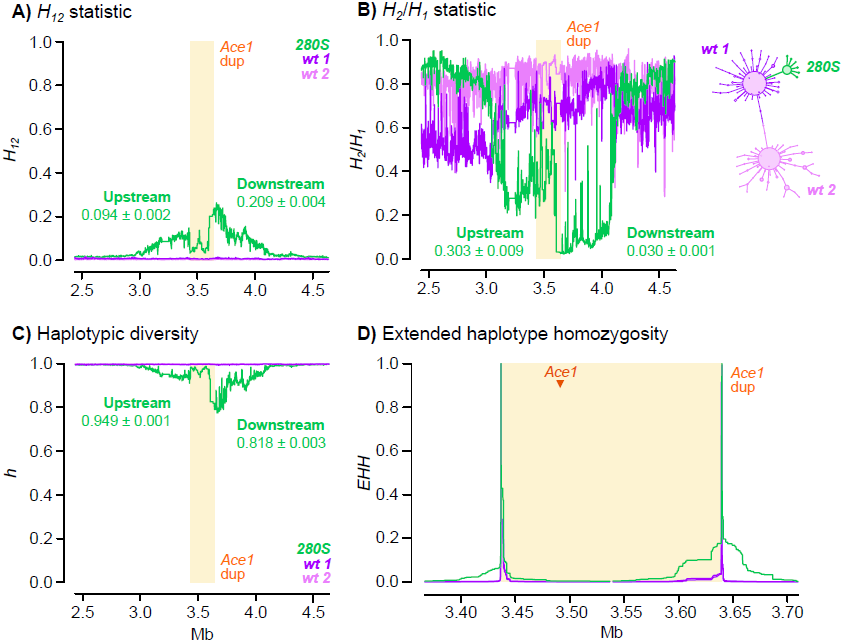
Positive selection around the *Ace1* duplication. **A-C)** Profile of Garud’s *H*_*12*_, *H*_*2*_*/H*_*1*_ and haplotypic diversity around the *Ace1* duplication, for haplotypes carrying the *280S*- or *wt*-tagging variants at 2R:3481632. Statistics are calculated in blocks of 500 variants with 20% block overlap. Includes averages of each statistic upstream and downstream of the breakpoints (with standard errors from jackknife haplotype resampling). **D)** Extended haplotype homozygosity at the duplication breakpoints for haplotypes carrying the *280S*- or *wt*-tagging variants at the 2R:3481632 locus. Additional plots for all tagging variants are available in Supplementary Material SM11 and SM12.

Next, we tested whether *280S* alleles and the duplication spread jointly between *A. coluzzii* and *A. gambiae* via introgression (Figure 8). We examined the chromosome-wide profile of introgression between *A. gambiae* and *A. coluzzii* populations using Patterson’s *D* statistic (Durand et al. 2011; Patterson et al. 2012), which compares allele frequencies between three putatively admixing populations (A, B and C) and one outgroup (O); and can identify introgression between populations A and C (*D* > 0) or B and C (*D* < 0). Specifically, we tested whether duplicated *A. coluzzii* from West African populations (population A) introgressed with *A. gambiae* populations from West and Central Africa (C), using *wt* Angolan *A. coluzzii* as a control (B) and *A. arabiensis* as an outgroup (O).

**Figure 8.**
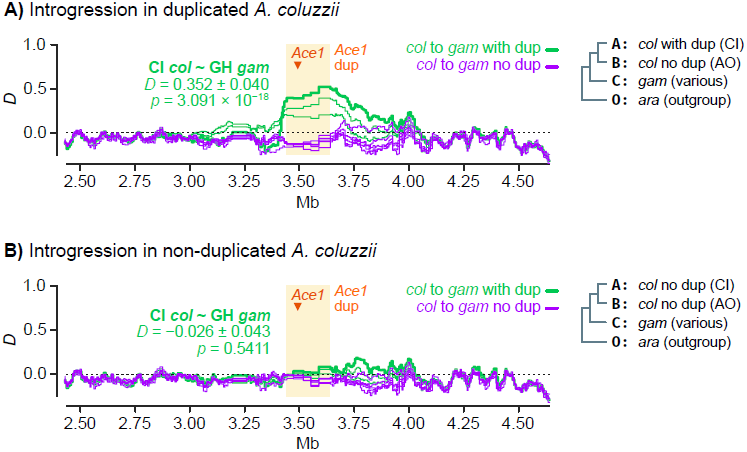
Introgression of the *Ace1* duplication. **A)** Profile of Patterson’s *D* statistic around the *Ace1* duplication, testing evidence of introgression between duplicated *A. coluzzii* from Côte d’Ivoire (population A) and various *A. gambiae* populations (population C) with or without duplications (green and purple lines, respectively). We used non-duplicated Angolan *A. coluzzii* as a contrast (population B) and *A. arabiensis* as outgroup (O). *D* is calculated in windows of 5,000 variants with 20% overlap, and the average value of D along the duplication region is shown for the Côte d’Ivoire *A. coluzzii* / Ghana *A. gambiae* comparison, with standard errors derived from block-jackknife. **B)** Id., but using non-duplicated *A. coluzzii* from Côte d’Ivoire as population A. Detailed lists of all statistical tests and replicate analyses with additional populations are available in Supplementary SM13. Species abbreviations: *gam, A. gambiae*; *col, A. coluzzii*; *ara, A. arabiensis*. Country abbreviations: AO, Angola; CI, Côte d’Ivoire; GH, Ghana.

We evaluated whether the *Ace1* duplication introgressed between *A. gambiae* and *A. coluzzii* in two scenarios: (i) in specimens with CNVs from populations with *Ace1* resistance mutations (Figure 8A), and (ii) in non-duplicated specimens from the same populations (Figure 8B). We only find evidence of introgression in the first case, i.e. between West African *A. gambiae* and *A. coluzzii* specimens carrying *Ace1* duplications (Figure 8A). For example, we found evidence of introgression between duplicated Ghanaian *A. gambiae* and duplicated Ivorian *A. coluzzii* (*D* = 0.352 +/- 0.040, *p* = 3.1 × 10^−18^ from a standard distribution of block-jackknifed estimates; green lines in Figure 8A); but not with non-duplicated Ivorian *A. coluzzii* (*D* = - 0.026 +/- 0.043, *p* = 0.54; green lines in Figure 8B). In all comparisons where introgression was apparent, the genomic region of increased *D* values extended along the entire duplicated region. On the other hand, there was no evidence of introgression in any comparison involving non-duplicated *A. gambiae* (*D* ≈ 0; purple lines in Figure 8A and 8B).

To establish the direction of introgression, we performed a phylogenomic analysis of haplotypes from the duplicated region (Figure 9). Firstly, haplotypes with *Ace1* duplications formed a single clade containing *A. gambiae* and *A. coluzzii* sequences to the exclusion of all non-duplicated sequences from both species. Secondly, we found that duplicated sequences were phylogenetically closer to the *wt A. gambiae* clade than to *wt A. coluzzii* (Figure 9A). Thirdly, an analysis of differences in allelic frequencies between *wt A. gambiae, wt A. coluzzii* and duplicated haplotypes also indicated that the duplicated haplotypes are more similar to *wt A. gambiae* than to *wt A. coluzzii*: the branch leading to *wt A. coluzzii* since the divergence from the duplicated specimens was longer (*L* = 0.0273 ± 0.0015 standard error) than the branch leading to *wt A. gambiae* (*L* = 0.0102 ± 0.0011). This topology indicates that duplicated haplotypes emerged from a *A. gambiae wt* background and later introgressed into *A. coluzzii*. In contrast, phylogenies built from non-introgressed regions upstream and downstream of the duplication exhibited a topology with species-specific clades that included both duplicated and non-duplicated specimens (Figure 9B, C).

**Figure 9.**
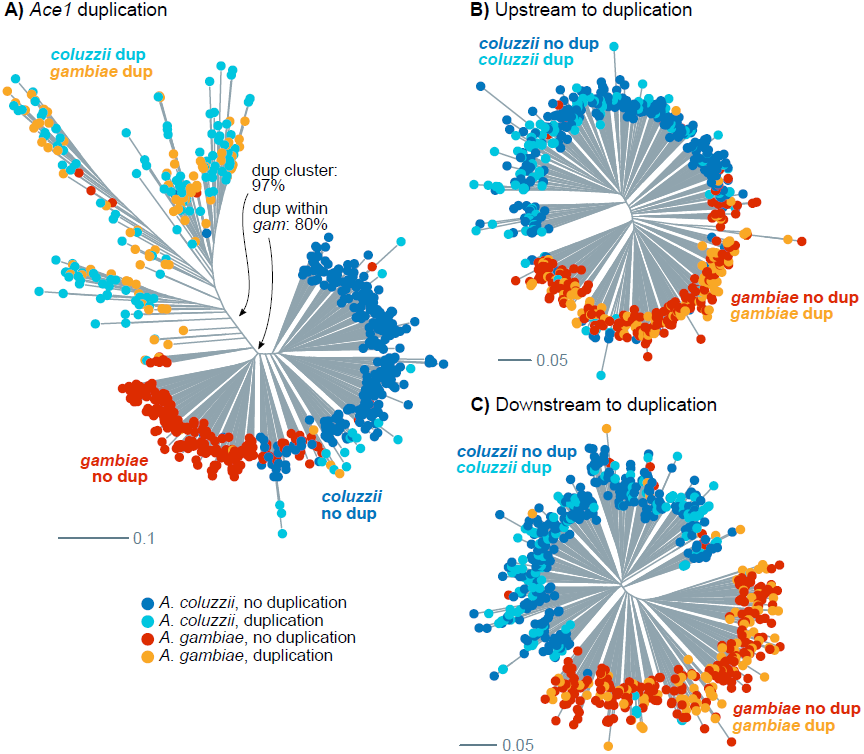
Phylogenetic analysis of introgression in *Ace1*. **A)** Maximum-Likelihood phylogenetic analysis of 690 Western African haplotypes from the *Ace1* duplicated region (2,787 phased variants), using a GTR model with ascertainment bias correction, empirical state frequencies and four Γ rate categories. **B-C)** Id., using variants upstream and downstream to the *Ace1* duplication (3,935 and 3,302 variants). Tips are color-coded according to species and duplication presence. Source alignments and complete phylogenies with supports in all nodes are available in Supplementary Material SM14 and SM15.

Altogether, these results show that the *280S* resistance alleles present across West Africa appear in similar genetic backgrounds irrespectively of the sampled population and species, and that this similarity extends beyond the *Ace1* gene to encompass the entire duplicated region (Figure 6). This low haplotypic diversity is due to a hard selective sweep, detectable at the duplication breakpoints (Figure 7). This resistance haplotype, which spans *ca*. 200 kbp and includes *Ace1* and ten other genes (Supplementary Material SM3) emerged in *A. gambiae* and later spread to *A. coluzzii* (Figures 8 and 9).

## Discussion

### Evolutionary history of *G280S* and *Ace1* duplications

Our results show that pirimiphos-methyl resistance is associated with a combination of two mutations in *Ace1*: the *G280S* SNP and CNVs. In West Africa, virtually all CNVs are found as duplications of a wide region (*ca*. 200 kbp) that includes *Ace1* and 10 other genes (Assogba et al. 2016; Lucas et al. 2019). This duplication has a unique evolutionary origin in the *A. gambiae*/*A. coluzzii* species pair, as its breakpoints are consistent in all populations from both species studied so far (Assogba et al. 2016; Lucas et al. 2019). The distribution of duplication-*280S* resistance haplotypes observed in the 1000 Genomes cohort can explained by three evolutionary events: the *G280S* mutation, an internally heterozygous duplication, and inter-specific introgression. Furthermore, recent sampling efforts have identified internal deletions within the duplication in both *A. gambiae* and *A. coluzzii* (Assogba et al. 2018). Homogeneous duplications are rare in the 1000 Genomes dataset but appear to be quite common in some *A. gambiae* populations, though less so in *A. coluzzii* (Table 2; (Weetman et al. 2015; Assogba et al. 2018)).

At present, we cannot establish the relative order of occurrence of the SNP and CNV in *Ace1* because these mutations are tightly linked in the 1000 Genomes dataset (Figure 2A). Nevertheless, the detection of *wt* homozygous duplications in samples collected in 2002 in southern Ghana (Weetman et al. 2015)—a country where *280S* alleles have more recently been found at high frequencies (Figure 1)—raises the possibility that the SNP might have occurred on an already duplicated background. This order of events would have initially resulted in permanent heterozygosity, providing resistance and reducing the fitness costs associated with impaired acetylcholinesterase activity in the absence of insecticide (Labbé et al. 2007; Assogba et al. 2015). If that were the case, the emergence of *280S-*homozygous duplications would require either a secondary loss of a *wt* copy or a parallel *G280S* mutation in the initially heterozygous duplication.

After these two mutations, the joint duplication-*280S* resistance haplotype introgressed from *A. gambiae* into *A. coluzzii* (Figures 8 and 9). This *A. gambiae* origin is consistent with earlier studies of *Ace1* variation in West African locations, which generally reported higher frequencies of *280S* (Djogbénou et al. 2008; Dabiré et al. 2009; Essandoh et al. 2013) and CNVs (Assogba et al. 2018) in *A. gambiae* than in *A. coluzzii*, as one would expect if they had a longer evolutionary history in the former. Whilst previous studies had suggested that the similarity of *280S* haplotypes in *A. coluzzii* and *A. gambiae* was due to inter-specific introgression (Djogbénou et al. 2008; Essandoh et al. 2013; Weetman et al. 2015), they were focused on partial sequencing of the *Ace1* gene and could not assess the relationship between introgression and the wider duplicated region (*ca*. 30 times longer than the *Ace1* coding region). Furthermore, analyses focused on the *Ace1* gene body can be confounded by the lack of sequence variation in this region (Supplementary SM9). By leveraging variation in linkage disequilibrium beyond the gene, we are able identify clear signatures of introgression along the entire *Ace1* duplication region (Figure 8).

### Genetic basis of pirimiphos-methyl resistance in *A. coluzzii* and *A. gambiae*

We have uncovered a strong association between resistance to pirimiphos-methyl and the possession of two or more copies of the *Ace1 280S* allele in *A. coluzzii* from Côte d’Ivoire (Figure 3). While the *G280S* mutation alone has been previously linked to resistance to carbamates and organophosphates in multiple insects (Weill et al. 2003, 2004; Labbé et al. 2007; Alout et al. 2007; Djogbénou et al. 2008; Dabiré et al. 2014; Liebman et al. 2015; Feyereisen et al. 2015), our results indicate that resistance to pirimiphos-methyl specifically relies on the presence of multiple *280S* alleles.

Across the 1000 Genomes cohort, specimens with *280S* alleles can be grouped into two categories according to the number of *Ace1* copies (Figure 2C): (i) heterozygous duplications with multiple *280S* alleles and/or multiple *wt*, and (ii) high-copy, *280S*-homozygous specimens. Heterozygous duplications are the most frequent combination of *Ace1* mutations (113 out of 119 samples, Figure 2B) and the only one identified in the *A. coluzzii* population from Côte d’Ivoire (Tiassalé). Therefore, we decided to examine their contribution to pirimiphos-methyl resistance in a wider array of *A. gambiae* and *A. coluzzii* populations from Ghana, Côte d’Ivoire, and Togo (Tables 1 and 2). We identified a common theme in most populations surveyed here: (i) *280S* heterozygotes were the most common resistance genotype in six out of eight populations analysed, including all *A. coluzzii* sampling locations and three out of five in *A. gambiae* (Table 1); and (ii) among the heterozygotes, we found that the ratio of *280S* alleles and *Ace1* copy number were significantly associated with resistance in five out of seven tested populations (Table 2). This provides a wider demonstration that the combined effect of *G280S* and *Ace1* duplications is significantly associated with pirimiphos-methyl resistance across multiple West African populations of both species; consistent with a requirement for multiple copies of *280S* for resistance.

We also identified six high-copy, *280S*-homozygous genomes sampled from the Ghanaian *A. gambiae* population in the 1000 Genomes dataset (Figures 1 and 2). These samples had not been assayed for pirimiphos-methyl resistance prior to sequencing. However, our extended genotype-phenotype analysis in eight West African populations (Table 1) showed that (i) *280S* homozygotes exhibited increased resistance to pirimiphos-methyl, and that, in fact, they were the most abundant resistance allele in two *A. gambiae* populations (Baguida and Obuasi). It is worth mentioning that survival rates among *280S* homozygotes were noticeably lower these two locations (56.8% - 58.9%) than in populations where heterozygotes were more common (> 95%; Table 1). Our analyses cannot fully capture the resistance landscape in these populations, because we have focused on the role of CNVs in heterozygous specimens (as they were the most abundant resistant type in the 1000 Genomes cohort; Figure 1). Thus, we cannot presently explain these differences, which may reflect an effect of different phenotyping strategies employed in each population (see Methods), or additional resistance mechanisms. A detailed examination of *280S*-homozygous duplications—e.g. effects on gene dosage or fitness—could shed light onto the phenotypic variability observed in populations with high penetrance of *Ace1* mutations.

Finally, we have performed a genome-wide scan in a population of *A. coluzzii* from Côte d’Ivoire (Tiassalé) to identify loci associated with pirimiphos-methyl resistance (Figure 5). Our investigation of the association between *k*-mer frequencies and resistance confirmed the overwhelming concentration of phenotypic association in the *Ace1* duplication (Figure 5B), supporting the conclusion that it is the primary driver of resistance in this population. The correlations between *Ace1* duplication and all but one of the *k*-mers significantly associated with resistance, but mapping elsewhere in the genome strongly suggests non-independence of the signals, but the exact cause of this is unclear. Whilst we cannot entirely discount linkage disequilibrium arising from epistasis, a simpler and parsimonious explanation would be that this result is caused by non-specific or incorrect sequence alignments, possibly linked to currently unrecognised variation in the *Ace1* duplication.

The lack of clear signals of pirimiphos-methyl adaptation other than *Ace1* in this population may reflect its collection at an early stage of the development of pirimiphos-methyl resistance in West Africa (Edi et al. 2014b). Therefore, our genome-wide analyses would only capture pre-existing variants that were enriched in the resistant specimens. *Ace1* mutations and duplications, which cause cross-resistance with previously employed insecticides (Djogbénou et al. 2015, 2008; Essandoh et al. 2013; Edi et al. 2014a; Assogba et al. 2015, 2016, 2018; Weetman et al. 2015), fit this definition and are sufficient to sustain high levels of resistance. However, future analyses on recent collections from populations subjected to regular pirimiphos-methyl-based IRS may reveal additional mechanisms more specifically selected by this insecticide.

### Implications for insecticide intervention programmes

Our results demonstrate that the duplication-*280S* haplotype represents a powerful marker to diagnose resistance to pirimiphos-methyl in *A. coluzzii* and *A. gambiae*, permitting the spread of resistance to be tracked from any preserved sample from which DNA can be extracted. Our initial survey of an *A. coluzzii* population from Tiassalé (Côte d’Ivoire) shows that discrimination between susceptible and resistant specimens can be attained with high accuracy in this population according to the number of *280S* alleles. Specifically, following the resistance thresholds apparent in Figure 3B, we can (i) classify *wt* homozygous samples as susceptible (100% predictive value) and (ii) separate heterozygous individuals between those with more *wt* than *280S* alleles (susceptible) and those with equal or more *280S* than *wt* alleles (resistant). This would yield a positive predictive value of 77% for resistance, and a negative predictive value of 85% for susceptibility.

In practice, precise quantification of the number of resistance *280S* alleles will often be difficult. Nevertheless, we show that key variation in *Ace1* can be captured using two qPCR genotyping assays that predict pirimiphos-methyl resistance in populations of *A. gambiae* and *A. coluzzii* (Table 2): (i) a measurement of the ratio of *280S* to *280G* alleles (Bass et al. 2010), and (ii) the number of *Ace1* copies (Edi et al. 2014a). Crucially, measurements of the *280S* allelic ratio can be obtained from the same assay that is currently used for standard genotyping of the *G280S* mutation (Bass et al. 2010), which thus has good diagnostic capacity in homozygotic and heterozygotic specimens. The phenotypic assays we employed were intended to provide more accurate separation than standard dose approaches (Weetman and Donnelly 2015), but the strong effect of *Ace1* CNVs and *G280S* mutations on pirimiphos-methyl resistance is sufficiently robust for resistance diagnosis in single time-dose discriminating assays as well – as used in the Ivorian *A. coluzzii* specimens from the 1000 Genomes cohort.

Furthermore, monitoring studies must note that this resistance haplotype is still evolving, especially in the light of the fitness trade-offs associated with *Ace1* resistance alleles – namely, that heterozygous duplications offset the fitness cost of *280S* alleles in the absence of insecticide exposure (Labbé et al. 2007; Assogba et al. 2015), and that variably copy number can result in gene dosage disturbances (Assogba et al. 2016). A recent survey has shown that internal deletions within the duplicated region, downstream to *Ace1*, are spreading in both *A. gambiae* and *A. coluzzii* (Assogba et al. 2018). The authors have proposed that these deletions reduce the fitness costs of duplications by ameliorating deleterious changes in gene expression (Assogba et al. 2018). These small deletions are absent from the Ivorian *A. coluzzii* genomes analysed here (Lucas et al. 2019), but future genome monitoring studies could investigate their spread and identify signals of selection associated with their proposed selective advantage. Likewise, the independently evolved *Ace1* CNV that we have identified in one *A. gambiae* specimen from Guinea (Figure 2B, Supplementary Material SM4) might merit further attention in the future. Finally, as noted above, we cannot rule out the possibility that, following several years of widespread use of pirimiphos-methyl for IRS, additional non-target site-based resistance mechanisms might have emerged.

The duplication-*280S* haplotype pre-dates the widespread adoption of pirimiphos-methyl in 2013 (Oxborough 2016), which indicates that it likely spread due to its adaptive value with respect to previously employed insecticides used in disease control or agriculture, such as carbamates (bendiocarb (Edi et al. 2014a; Assogba et al. 2016)) or other organophosphates (fenitrothion, chlorpyrifos-methyl (Essandoh et al. 2013; Assogba et al. 2016)). In that regard, we show that the evolution of the pirimiphos-methyl resistance haplotype is deeply intertwined with that of previous insecticide adaptations, and reveal how continued genomic monitoring studies can help us understand the influence of previous interventions on future population control efforts. Furthermore, whilst current efficacy of pirimiphos-methyl in Actellic IRS formulations appears to be retained (Sherrard-Smith et al. 2018), its potential could be limited in the *Anopheles* populations where resistance to these insecticides is already common, most notably in West Africa (Figures 1 and 2). This emphasises the crucial importance of now-commencing resistance management strategies using additional IRS insecticides with alternative modes of action (Oxborough et al. 2019).

## Materials and methods

### Data collection

We used genomic variation data from individual mosquitoes, obtained from the Phase 2 (AR1) of the *Anopheles gambiae* 1000 Genomes project, as described in (Miles et al. 2017; The *Anopheles gambiae* 1000 Genomes Consortium 2019). This dataset consists of 1,142 wild-caught mosquitoes (1,058 females and 84 males) from 33 sampling sites located in 13 sub-Saharan African countries (listed in Supplementary Material SM4), and collected at different times between 2009 and 2012 (with the exception of samples from Gabon and Equatorial Guinea, collected in 2000 and 2002 respectively).

A detailed explanation of the methods used in whole-genome sequencing and variant calling can be found in the original publications (Miles et al. 2017; The *Anopheles gambiae* 1000 Genomes Consortium 2019). Briefly, DNA from each specimen was extracted with a Qiagen DNeasy Blood and Tissue Kit (Qiagen Science, US) and sequenced with the Illumina HiSeq 2000 platform (Wellcome Sanger Institute, UK) using paired-end libraries (100 bp reads, 100-200 bp insert sizes), aiming at a 30× coverage per specimen. Variant calling was performed using *bwa* (Li and Durbin 2009) 0.6.2 and the *GATK* 2.7.4 *UnifiedGenotyper* (Van der Auwera et al. 2013). Haplotype phasing was estimated with *SHAPEIT2* (Delaneau et al. 2013), and variant effects were predicted using *SnpEff* 4.1b (Cingolani et al. 2012). We downloaded the genotype calls, SNP effect predictions, and haplotype phasing as available in *Anopheles gambiae* 1000 Genomes Phase 2 online archive (The *Anopheles gambiae* 1000 Genomes Consortium 2017).

The catalog of *Ace1* CNVs in the 1000 Genomes dataset (Phase 2) was obtained from (Lucas et al. 2019). There, the authors used calculated the coverage (sequencing depth) of each whole-genome sequenced sample in non-overlapping windows along the genome (300 bp) and normalised this value so as to obtain an expected average value of coverage = 2 in non-duplicated, diploid regions. A Gaussian HMM procedure was then applied to the normalised windowed coverage data so as to call windows with heightened normalised coverage (>2). A detailed account of these methods can be found in the original publication (Lucas et al. 2019), and the results for the *Ace1* region are available in Supplementary Material SM4.

We retrieved the reference genome information for *A. gambiae* from VectorBase (Giraldo-Calderón et al. 2015), including the genome assembly (PEST, AgamP4 version), gene annotation coordinates in GFF format (AgamP4.9) and gene names and descriptions.

### Genotype-phenotype association in Côte d’Ivoire *A. coluzzii* genomes

As part of *Anopheles gambiae* 1000 Genomes Phase 2, 71 *A. coluzzii* specimens from Côte d’Ivoire were collected and phenotyped for pirimiphos-methyl resistance before whole-genome sequencing (The *Anopheles gambiae* 1000 Genomes Consortium 2019). These samples were collected as larvae in rice fields near Tiassalé (coordinates: -4.823, 5.898) between May and September 2012, and were identified as *A. coluzzii* based on PCR assay (Fanello et al. 2002). The 71 larvae were tested for resistance to pirimiphos-methyl resistance using a WHO tube assay with 0.25% impregnated papers (World Health Organization 2018b), which led to the identification of 31 resistant (live) and 40 susceptible (dead) specimens.

We tested the association of these resistance phenotypes with various genetic variants present in *Ace1* in this population. This includes the non-synonymous mutations *G280S* and *S65A*, the number of *Ace1* copies, and the number of *280S* alleles in each individual sample (listed in Supplementary Material SM6A).

For the non-synonymous mutations, we assessed genotype-phenotype associations for each variant independently using Fisher’s exact test (*gmodels R* library (Warnes et al. 2018)) and estimated odds ratio and 95% confidence intervals using the Woolf method with Haldane-Anscombe correction (*Prop*.*or* function in the *pairwiseCI R* library (Schaarschmidt and Gerhard 2019)). For the *Ace1* and *280S* copy numbers, we used used binomial generalised linear models (*glm* function in *R stats* library, family *binomial*) to obtain odds ratio estimates for each of the four variables.

We also built a binomial generalised linear models with all four variables (*G280S* and *A65S* genotypes; *Ace1* and *280S* copy number) and a step-wise procedure to remove non-informative variants from the model (*step* function in *R stats* using *k = log(n)* as the degree of freedom, as required by the Bayesian Information Criterion; *n* represents the number of samples in our dataset). The number of *Ace1* copies and the number of mutated alleles were encoded as continuous variables, and the genotypes of *G280S* and *S65A* were encoded as categorical variables. The performance of all generalised linear models was assessed relative to a null model (intercept as the only variable) using a *χ*^*2*^ test in an analysis of variance (*anova* function in the *R stats* library). A detailed statistical analysis of all comparisons mentioned above can be found in Supplementary Material SM6A and SM6B.

The number of *Ace1* copies in each genome of *Anopheles gambiae* 1000 Genomes Phase 2 was retrieved from (Lucas et al. 2019). In that study, copy number states of multiple CNVs were inferred using a HMM-based predictive model that took as input the normalised sequencing depth calculated along non-overlapping 300 bp genomic windows.

The estimated number of *280S* alleles in each genome was estimated in the following manner: (i) we retrieved the copy numbers of *Ace1* in each genome (*C*; see above); (ii) we calculated the fraction of reads supporting the reference and alternate alleles; and (iii) assigned the number of *280S* alleles (*S*) as the value that minimised the difference between the fraction of alternate alleles and *S/C*, for all discrete *S* values between 0 and *C*. For example, a genome with three *Ace1* copies (*C* = 3) and 30% of reads supporting *280S* would carry one *280S* allele (*S* = 1), given that *S/C* = 1/3 ≈ 30%.

### Genotype-phenotype association in additional West African populations

We collected 1080 female specimens of two species (*A. coluzzii* and *A. gambiae*) from six different locations across West Africa (Côte d’Ivoire: Aboisso; Ghana: Madina, Korle-Bu, Weija and Obuasi; Togo: Baguida). Species identity was determined using two methods designed to discriminate between *A. gambiae, A. coluzzii* and *A. arabiensis*: a PCR of species-specific SINE insertion polymorphisms as described in (Santolamazza et al. 2008), and a high-resolution melt curve analysis (Chabi et al. 2019). Details for each of these methods, including primer sequences, are available in Supplementary Material SM6G. The complete list of specimens, sampling times and locations, and species identification are available in Supplementary Material SM6C and SM6E.

All 1080 specimens were phenotyped for pirimiphos-methyl resistance using a custom dose-response assay with WHO standard tubes (World Health Organization 2018b). For the Aboisso (Côte d’Ivoire), Korle-Bu, Weija and Madina samples, resistant specimens were determined as surviving a threshold concentration of pirimiphos-methyl after exposure during the larval stage, and susceptible specimens were identified as dead after a lower threshold dose (constant exposure time of 60 minutes). The exact concentration thresholds for the dose-response assay were determined on a per-site basis, and are listed in Supplementary Material SM6C. For example, *A. gambiae* from Aboisso were deemed resistant if they survived a 1x concentration of pirimiphos-methyl (0.25%) for 60 minutes, and susceptible if they died at 0.5x concentration (0.125%). In the Obuasi (Ghana) and Baguida (Togo) populations, phenotypes were determined according to survival after different exposure times (susceptible: dead after less than 45 minutes; resistant: alive after 45 minutes or more) using a single concentration of pirimiphos-methyl (0.5x, 0.125%). The exact exposure time for each sample in intervals of 15 minutes are listed in Supplementary Material SM6C.

Out of 1080 specimens, 909 could be assigned to one of the following populations, defined as combinations of species and collection locations with determined resistance phenotypes: *A. coluzzii* from Aboisso, Côte d’Ivoire (*n =* 55); *A. coluzzii* from Korle-Bu (*n* = 214), and Weija (*n* = 131), Ghana; *A. gambiae* from Aboisso, Côte d’Ivoire (*n =* 82); *A. gambiae* from Baguida, Togo (*n* = 102); *A. gambiae* from Madina (*n* = 172), Obuasi (*n* = 140) and Weija (*n* = 13), Ghana.

For each sample, we determined the *G280S* genotype using a qPCR TaqMan assay as described by Bass *et al*. (Bass et al. 2010) (list of samples in Supplementary Material SM6C), in which *280S*- and *280G*-specific primers where labeled with different fluorescent dyes (FAM and HEX, respectively). We also calculated the ratio of FAM-to-HEX fluorescent dye signal in hetrerozygotes, which label *280S* and *280G* alleles respectively, as an index of the fraction of *280S* allele copies (Edi et al. 2014a; Djogbénou et al. 2015). Detailed methods for these two genotyping assays are available in Supplementary Material SM6G.

We assessed the association of *G280S* mutations with resistance for each of the populations listed above using generalised linear models (*glm* function in *R stats* library, *binomial* family). The results from these tests are available in Supplementary Material SM6D.

A sub-set of these 909 specimens (*n* = 167; listed in Supplementary Material SM6E) was also genotyped for CNV polymorphisms in the *Ace1* locus using a qPCR assay and a combination of primers for *Ace1* and two non-duplicated control genes (the *CYP4G16* AGAP001076; and the elongation factor AGAP005128). Detailed methods for CNV genotyping assay are available in Supplementary Material SM6G and (Edi et al. 2014a). These samples were selected from the subset of heterozygotes in order to investigate the effect of both CNVs and fraction of *280S* alleles on resistance.

We used generalised linear models (*glm* function in *R stats* library, *binomial* family) to assess the effect of these two variables within each of the seven populations with available CNV data (three from *A. coluzzii*, four from *A. gambiae*; listed in Table 2 and Supplementary Material SM6F). Then, we obtained the minimal significant model for each population using a stepwise reduction procedure and the BIC criterion (*step* function in the *R stats* library), and assessed their fit against a null model (intercept as the only variable) with an analysis of variance and a *χ*^*2*^ test (*anova* function in *R stats*). The results from these tests are available in Supplementary Material SM6F.

### Haplotype clustering and analysis of selection signals in *Ace1*

Haplotypes along chromosome arm 2R were reconstructed using the dataset of phased variants from the *Anopheles gambiae* 1000 Genomes Phase 2 data. Specifically, we retained phased variants that were biallelic, non-singletons and segregating in at least one population.

Given the paucity of phased variants within the *Ace1* gene body (Supplementary SM9A, B) and the associated difficulties in phasing the *G280S* mutation itself, we analysed the haplotypes around nearby genetically linked variants instead. To do so, we calculated the linkage disequilibrium between position *G280S* and all variants located between positions 3416800 and 3659600 along 2R (i.e., 20kbp upstream and downstream of the duplication breakpoints [3436800-3639600]). Linkage disequilibrium was estimated from genotype counts in each sample using Rogers’ and Huff *r* correlation statistic (Rogers and Huff 2009) as implemented in *scikit-allel* v1.1.10 library (*rogers_huff_r*) (Miles and Harding 2017). Then, we retrieved all variants with *r* > 0.95 that were also present in the subset of phased variants (four in total). These four linked variants had minor allele frequencies similar to that of *280S* in Côte d’Ivoire (43%) and were located in intergenic regions −26 kbp and +12 kbp of *G280S* (listed in Supplementary Material SM9C).

We retrieved haplotypes surrounding each of the four tagging variants and clustered them by similarity using minimum spanning trees. Specifically, we retrieved phased variants ± 300 bp of each tagging variant (retaining variants that were biallelic, non-singletons and segregating in at least one population; these regions contained between 67 and 164 phased variants depending on the analysis; Supplementary Material SM10). We used allele presence/absence data from each haplotype to build minimum spanning trees of haplotypes based on Hamming distances (breaking edges for distances >1). This distance matrix was then used to build medium spanning trees (*minimum_spanning_tree* function in *SciPy* 1.1.0 Python library (Jones et al. 2019), from the *sparse*.*csgraph* submodule). Tree visualizations were produced using the *graphviz* 2.38.0 Python library (Ellson et al.) and the *hapclust* library (Clarkson et al. 2018; Clarkson and Miles 2018), and clusters of haplotypes were color-coded according to the population of origin or the presence/absence of the alternate allele (Supplementary Material SM10).

We used these trees to identify groups of highly similar haplotypes linked to the *280S* or *wt* alleles in *Ace1* (Supplementary Material SM11). We calculated the profile of Garud’s *H* statistics (*moving_garud_h* function, *scikit-allel*) (Garud et al. 2015; Messer and Petrov 2013) and haplotype diversity (*moving_haplotype_diversity* in *scikit-allel*) around the *Ace1* duplication region in windows of consecutive variants (500 variants with 20% overlap). These calculations were performed separately for the main groups of identical haplotypes (*280S-*linked or *wt-*linked) as identified around each of the tagging variants (minimum size = 100 haplotypes). In addition, we calculated the averages of these same statistics in the regions immediately upstream and downstream of the duplication breakpoints (2R:3436800 – 5000 bp and 2R:3639600 + 5000 bp, respectively), and we calculated standard errors of these estimates using sample jack-knifing.

Finally, we calculated the rates of extended haplotype homozygosity decay for each cluster (Supplementary Material SM12), using 10,000 phased variants upstream and downstream of the duplication breakpoints (*ehh_decay* function in *scikit-allel*).

### Introgression analysis

We used Patterson’s *D* statistic (Durand et al. 2011; Patterson et al. 2012) to detect introgression between *A. coluzzii* and *A. gambiae*. We retrieved the allele frequencies of all biallelic, non-singleton variants from chromosome arm 2R that were segregating in West African populations where we had identified *Ace1* duplications (namely: 75 *A. coluzzii* from Burkina Faso, 55 from Ghana, and 71 from Côte d’Ivoire; and 92 *A. gambiae* from Burkina Faso, 12 from Ghana and 40 from Guinea; total *n* = 345 genomes) (Lucas et al. 2019), as well as Central African populations that we used as non-admixed controls (78 *A. coluzzii* from Angola; 69 *A. gambiae* from Gabon and 297 from Cameroon). In addition, we retrieved the genotypes at the same variant positions for the populations of *A. arabiensis* (12 genomes), *A. quadriannulatus* (10), *A. merus* (10) and *A. melas* (4) populations analysed in (Fontaine et al. 2015), which we used as outgroups in the calculation of Patterson’s *D*. In total, we retained *ca*. 12% of the 47,817,813 variants in 2R in each analysis.

To determine whether duplicated and non-duplicated populations had different introgression patterns in *Ace1*, we calculated windowed means of Patterson’s *D* (length 5,000 variants, 20% step; *moving_patterson_d* in *scikit-allel*) using various combinations of populations with and without the *Ace1* duplication (Figure 5A). This test requires genotype frequencies in four populations (two ingroups A and B; one candidate donor/receptor C with which A or B might have introgressed; and one unadmixed outgroup O) branching in a (((A,B),C),O) topology (Figure 5A). We observed the following convention: (i) we used West African *A. coluzzii* populations as A, discriminating between duplicated and non-duplicated subpopulations; (ii) we used non-duplicated *A. coluzzii* from Angola as control B; (iii) we used various *A. gambiae* populations as C, again analysing duplicated and non-duplicated specimens within each population separately; and (iv) *A. arabiensis, A. quadriannulatus, A. merus* and *A. melas* as outgroup O.

For each combination of (((A,B),C),O) populations, we calculated the average *D* statistic within the duplicated (position 3,436,800 to 3,639,600, spanning 202.8 kbp around *Ace1*), and estimated its deviation from the null expectation (no introgression: *D* = 0) with a block-jackknife procedure (block length = 100 variants; *average_patterson_d* function in *scikit*-*allel*), which we used to estimate the standard error, *Z-*score and the corresponding *p* value from the two-sided normal distribution (Supplementary Material SM13).

In parallel, we performed a phylogenomic analysis of the reconstructed haplotypes of the *Ace1* CNV region (position 3436800 to 3639600) in the West African populations where duplications had been identified (75 *A. coluzzii* from Burkina Faso, 55 from Ghana, and 71 from Côte d’Ivoire; and 92 *A. gambiae* from Burkina Faso, 12 from Ghana and 40 from Guinea; total *n* = 345 genomes, 690 haplotypes). Specifically, we built an alignment of phased variants along the duplication (2,787 positions) and computed a Maximum-Likelihood phylogenetic tree using *IQ-TREE* 1.6.10 (Nguyen et al. 2015). We used the GTR nucleotide substitution model with ascertainment bias correction, empirical state frequencies observed from each alignment, and four gamma distribution (*Γ*) categories (*GTR+F+ASC+G4* model in *IQ-TREE*). This model was selected by the *IQ-TREE* implementation of *ModelFinder* (Kalyaanamoorthy et al. 2017) (*TEST* mode) as the best-fitting model according to the BIC criterion. The best-fitting tree was found after 22,000 iterations (correlation coefficient threshold = 0.99). We estimated node statistical supports from 1,000 UF bootstraps (Hoang et al. 2018; Minh et al. 2013). Using the same approach, we built two additional phylogenies from genomic regions located upstream and downstream of the duplication breakpoints (3,935 and 3,302 variants extracted from 50 kbp segments located −1Mb and +1Mb), finding the best-fitting trees after 7,659 and 27,400 iterations, respectively (correlation coefficient threshold = 0.99). The resulting phylogenetic trees were plotted, unrooted, using the *phytools* 0.6-60 (Revell 2012) and ape 5.3 libraries (*plot*.*phylo*) (Paradis and Schliep 2019). Phylogenies and source alignments are available as Supplementary Material SM14 and SM15.

We also used allelic frequencies within the inversion to estimate the divergence times between specimens carrying the duplication (both *A. gambiae* and *A. coluzzii*), *wt A. coluzzii* and *wt A. gambiae*. This three-way calculation of divergence times reflects the amount of allele frequency change between one of the three groups relative to the other two, and is therefore analogous to the population branch statistic (Cavalli-Sforza 1969; Yi et al. 2010). Following this logic, we estimated separately (i) the branch length of *wt A. coluzzii* relative to the separation of *wt A. gambiae* and the duplicated sequences, and (ii) the branch length of *wt A. gambiae* relative to the separation of *wt A. coluzzii* and the duplicated sequences. We performed these calculations in non-overlapping windows of 100 variants with the *pbs* function in *scikit-allel*, and estimated standard errors using a block-jackknifing procedure.

### Genetic differentiation in *A. coluzzii* from Côte d’Ivoire

We assessed the degree of genetic differentiation between resistant and susceptible *A. coluzzii* from Côte d’Ivoire. We focused on non-singleton variants that were segregating in this population. In total, 8,431,869 out of 57,837,885 (14.57%) variants from chromosomes 2, 3 and X were retained for further analysis.

For each chromosome arm (2R, 2L, 3R and X), we calculated genotype counts in each subpopulation (resistant and susceptible), and calculated their genetic differentiation using the Hudson’s *F*_*ST*_ statistic (Hudson et al. 1992; Bhatia et al. 2013) along non-overlapping windows of 1,000 variants (*moving_hudson_fst* function in the *scikit-allel* v1.1.10 library (Miles and Harding 2017) from Python 3.4). We also calculated the average *F*_*ST*_ in each chromosomal arm, with standard errors obtained from a block-jackknife procedure (using non-overlapping windows of 1,000 variants; *average_hudson_fst* function).

We calculated the normalised population branching statistic (*PBS*) (Yi et al. 2010; Malaspinas et al. 2016) in non-overlapping windows of 1,000 variants along each chromosomal arm (*pbs* function in *scikit-allel*). using resistant and susceptible Ivorian *A. coluzzii* as test populations, and *A. coluzzii* from Angola as an outgroup. Angolan *A. coluzzii* (*n* = 78) where selected as outgroup due to their relative isolation relative to West African *A. coluzzii* populations (The *Anopheles gambiae* 1000 Genomes Consortium 2019; Miles et al. 2017) and their putatively naive profile of organophosphate resistance (Oxborough 2016; World Health Organization 2018a). The distribution of *PBS* estimates along each chromosomal arm was standardised to unit variance (*standardize* function in *scikit-allel*), and the resulting distribution of *Z-*scores was used to derive a two-sided *p*-value that reflected extreme values of *PBS* (highly positive or negative). We corrected for multiple testing using local estimation of false discovery rates (*fdrtool* function in the *fdrtool* 1.2.15 *R* library (Klaus and Strimmer. 2015)). Finally, we selected genomic windows with high differentiation in resistant specimens (standardised *PBS* > 0, significance threshold *FDR* < 0.001).

The estimates of *PBS* and Hudson’s *F*_*ST*_ in each genomic window are available as Supplementary Material SM7.

We performed a principal component analysis of resistant and susceptible Ivorian *A. coluzzii*. We first obtained a set of genetically unlinked variants from chromosomal arms 3R and 3L (so as to avoid the confounding effects of chromosomal inversions in arms 2R, 2L and X). We discarded linked variants within 500 bp consecutive windows (using a 200 bp step), using a Rogers and Huff’s *r* threshold value of 0.1 (*locate_unlinked* function in *scikit-allel*), and repeated this process for ten iterations for each chromosome arm. This filtering procedure resulted in 791 genetically unlinked variants from both chromosomal arms. We used the alternate allele count in each variant to construct a PCA using singular value decomposition (*pca* function from *scikit-allel*), and scaling the resulting coordinates using Patterson’s procedure (Patterson et al. 2006).

### *k*-mer enrichment analysis in *A. coluzzii* from Côte d’Ivoire

We obtained *k*-mer counts for each of the 71 *A. coluzzii* samples using the *count* function in *jellyfish* v. 2.2.10 (Marçais and Kingsford 2011), using a *k* = 31 bp (parameters: *-C -m 31 --out-counter-len 2 -s 300M --bf-size 10G*). To reduce the computer memory footprint of the *k*-mer count tables, the *k*-mer strings were recoded as integers, split into separate lexicographical groups according to the leading nucleotides (i.e. *k*-mers beginning with *AAA, AAC*, etc. were assigned to different groups), and, within each set, sorted lexicographically again. In total, we recorded the frequency of 1,734,834,987 *k*-mers across 71 samples. The resulting count tables were filtered to retain *k*-mers showing variation in the population: we discarded *k*-mers present in fewer than 3 samples, or absent in fewer than 3 samples. These filters removed 967,274,879 *k*-mers, leaving 767,560,108 *k*-mers for further analysis.

To test whether the frequencies of any *k*-mers were associated with pirimiphos-methyl resistance, normalised *k*-mer frequencies were obtained by dividing the *k*-mer counts by the total number of variant *k*-mers in each sample. We calculated the Spearman’s rank correlation of each *k*-mer frequency with the resistance phenotype for each of the 767,560,108 *k*-mers (*cor*.*test* function in the *R stats* library (R Core Team 2017)), correcting for multiple testing using local estimation of false discovery rates (*fdrtool* function in *fdrtool* library (Klaus and Strimmer. 2015)), and using a significance threshold of *FDR* < 0.001.

Given that multiple *k*-mers can overlap a single mutation, it was likely that many of the *k*-mers identified as significant were overlapping. We therefore took the *k*-mers that showed a significant association with pirimiphos-methyl resistance and assembled them by joining any *k*-mers that overlapped perfectly over at least 10 bp. The resulting assembled *k*-mers were aligned against the *A. gambiae* reference genome using *bwa mem* version 0.7.12-r1034 (parameters: *-T 0*) (Li and Durbin 2009). The mapping coordinates and sequences of the significant assembled *k*-mers are available in Supplementary Material SM8.

A minority of assembled *k*-mers aligned in regions other than the *Ace1* duplication. To determine whether this was due to mis-alignment, we correlated the frequency of these assembled *k*-mers in each sample against the *Ace1* copy number. Frequency for each assembled *k*-mer was calculated as the mean normalised frequency of all the *k*-mers that were used in its assembly. Because these *k*-mers and the *Ace1* copy number share the property of being correlated with pirimiphos-methyl resistance phenotype, they are statistically likely to have correlated frequencies even if physically unlinked. To control for this, the residuals of *k*-mer frequency and *Ace1* copy number were calculated within each of the resistant and susceptible groups of samples. Pearson’s correlation was then calculated on these residuals for each of the assembled *k*-mers (available in Supplementary Material SM8C).

Scripts to reproduce the *k*-mer analysis steps described above (counting, filtering, significance testing, assembly, mapping, and *Ace1* correlation analysis) are available on Github (see Availability of data and materials).

### Alignment and phylogenetic analysis of ACE proteins in animals

To obtain a candidate list of homologs of the *Ace1* gene, we retrieved all genes belonging to the orthogroup 339133at33208 from OrthoDB (Kriventseva et al. 2019) (682 genes in total). We aligned these candidates to a curated database of predicted proteomes from 89 complete animal genomes (listed in Supplementary Material SM2, including data sources) using Diamond 0.9.22.123 (Buchfink et al. 2014), and retained all alignments with an identity >95% to any of the candidate queries (130 genes). Next, we performed a multi-sequence alignment with *MAFFT* 7.310 (1,000 rounds of iterative refinement, L-INS-i algorithm) (Katoh and Standley 2013) of these 130 genes, together with the truncated sequence of the Pacific electric ray *Torpedo californica* protein used for its crystal structure analysis (PDB accession number: 1W75) and the full-sequence of the ortholog from its close relative *T. marmorata* (Uniprot accession number: P07692). This alignment (n=132 genes) was trimmed position-wise using *trimAL* 1.4 (Capella-Gutiérrez et al. 2009) (*automated1* mode), and 277 conserved columns were retained. A maximum-likelihood phylogenetic analysis of the trimmed alignment was then performed using *IQ-TREE* 1.6.10 (Nguyen et al. 2015), using a LG substitution matrix (Le and Gascuel 2008) with four G categories and accounting for invariant sites (*LG+I+G4* model). *IQ-TREE* was run for 469 iterations until convergence was attained (correlation coefficient ≥ 0.99). Node statistical supports were calculated using the UF bootstrap procedure (1,000 replicates) (Hoang et al. 2018; Minh et al. 2013). Alignment visualizations (Supplementary Material SM1) were obtained from Geneious 11.1.4 (Geneious 2019). Complete alignments are available as Supplementary Material SM2.

## Availability of data and materials

*Python* 3.4.7 and *R* 3.6.2 scripts to reproduce the main analyses from the paper are available in the following Github repository: github.com/xgrau/ace1-anopheles-report.

All genome variation data has been obtained from the publicly available repositories of the *Anopheles gambiae* 1000 Genomes project (Phase 2 AR1) (The *Anopheles gambiae* 1000 Genomes Consortium 2017): https://www.malariagen.net/data/ag1000g-phase-2-ar1 (download instructions can be found in the Github repository). All quantitative PCR data are provided as Supplementary Material SM6.

## Supporting information

SM1. Homology of Ace1 mutations

SM2. Alignments of ACE homologs

SM3. Ace1 mutations

SM4. Samples from Ag1000G Phase 2

SM5. Genes in Ace1 duplication

SM6. Phenotype-genotype association analyses

SM7. Genetic differentiation in Ivorian A. coluzzii

SM8. k-mer analysis in Ivorian A. coluzzii

SM9. Identification of tagging variants for Ace1 280S

SM10. Haplotype networks around the Ace1 tagging variants

SM11. Signals of selection in the Ace1 duplication

SM12. Extended haplotype homozygosity in the Ace1 duplication

SM13. Introgression of the Ace1 duplication

SM14. Haplotype alignments

SM15. Haplotype phylogenies

## Acknowledgements

We thank Sean Tomlinson (LSTM) and Chris Clarkson (Wellcome Sanger Institute) for fruitful discussions on the analyses.

## Supplementary Material legends

**Supplementary Material SM1. Homology of *Ace1* mutations. A)** Alignment of ACE homologs (protein sequences) in selected species (*A. gambiae, Culex quinquefasciatus, Aedes aegypti, Homo sapiens*, and *Torpedo californica*), used to determine the homology of non-synonymous mutations in this gene (*A65S* and *G280S* are highlighted). **B)** Alignment of ACE protein homologs in 20 culicine species, focusing on the vicinity of codon 280 (highlighted). **C)** Maximum-Likelihood phylogenetic analysis of ACE homologs from 89 animals (listed in Supplementary Material SM2, including data sources and accession numbers).

**Supplementary Material SM2. Alignments of ACE homologs. A)** Peptide coordinates of *A. gambiae Ace1* codon 280 in orthologs of *Ace1* from 89 animal genomes. **B)** Alignment of *Ace1* homologs from 89 animal genomes. **C)** Data sources, accession numbers, species abbreviation and taxonomy of the 89 animal genomes used in the analysis of ACE homology.

**Supplementary Material SM3. *Ace1* mutations**. Coordinates of non-synonymous mutations in *Ace1* and frequencies in each population of the *Anopheles gambiae* 1000 Genomes Phase 2 dataset. Species codes: *A. gambiae*, gam, *A. coluzzii*, col. Population codes: Angola, AOcol; Burkina-Faso, BFcol and BFgam; Côte d’Ivoire, CIcol; Cameroon, CMgam; Mayotte, FRgam; Gabon, GAgam; Ghana, GHcol and GHgam; The Gambia, GM; Guinea, GNcol and GNgam; Equatorial Guinea, GQgam; Guinea-Bissau, GW; Kenya, KE; Uganda, UGgam.

**Supplementary Material SM4. Samples from *Ag1000G* Phase 2**. List of samples from the *Anopheles gambiae* 1000 Genomes Phase 2 dataset with accession numbers, sample metadata (population and region of origin, collection date, species, sex), and summary of the main *Ace1* mutations in each sample (genotypes in the *G280S* mutation, number of reads supporting *280G* and *280S* alleles, number of CNVs). CNV coordinates and copy number from Lucas *et al*. (2019).

**Supplementary Material SM5. Genes in *Ace1* duplication**. List of genes in the *Ace1* duplication region, with genomic coordinates along the 2R chromosomal arm.

**Supplementary Material SM6. Phenotype-genotype association analyses. A)** Genotypes of *G280S* and *A65S* mutations in *A. coluzzii* samples from Côte d’Ivoire (*Anopheles gambiae* 1000 Genomes Phase 2), and resistance phenotype to pirimiphos-methyl. **B)** Phenotype-genotype association tests for the 71 Ivorian *A. coluzzii* samples. Includes summaries of single-variable GLM models for the following variables: *280S* presence (including a subset of samples with only one *280S* allele), *65S*, CNV and number of *280S* alleles. Also includes the minimal model obtained using stepwise reduction of a starting multi-variable model (BIC criterion). Model significance is measured with ANOVA comparison with null model (no variables) and a χ^2^ test. **C)** *G280S* genotypes and pirimiphos-methyl resistance phenotype for 1080 mosquitoes from West African populations of *A. gambiae* and *A. coluzzii* collected from six locations. For each sample, we also report collection dates, geographical locations, and details of the species identification, genotyping, and phenotyping (concentrations, exposure time). **D)** Phenotype-genotype association tests for *G280S* alleles in eight West African populations. Includes summaries of GLM models for each population. Model significance is measured with ANOVA comparison with null model (no variables) and a χ^2^ test. **E)** Number of *Ace1* copies and pirimiphos-methyl resistance phenotypes for 167 mosquitoes from the same West African populations. **F)** Phenotype-genotype association tests for CNV mutations (*Ace1* copies [CNV] and *280S-*to-*280G* allele ratio [ratio_FAM_HEX]) in West African populations. Includes summaries of GLM models for each population and each variable separately, and a minimal GLM obtained using the BIC criterion from an initial model with both variables. Model significance is measured with ANOVA comparison with null model (no variables) and a χ^2^ test. **G)** Protocols for species identification and *Ace1* mutation genotyping, including primer sequences.

**Supplementary Material SM7. Genetic differentiation in Ivorian *A. coluzzii*. A)** Genetic differentiation statistics (Hudson’s *F*_*ST*_ and *PBS*) between resistant and susceptible *A. coluzzii* from Côte d’Ivoire, in windows of 10,000 variants along the genome. *PBS* values are calculated using Angolan *A. coluzzii* as outgroup, and we also report a *Z*-score and *p*-value derived from a two-sided normal distribution. **B)** Genes overlapping the regions of high *PBS* (*FDR* < 0.001 and standardised *PBS* > 0).

**Supplementary Material SM8. *k*-mer analysis in Ivorian *A. coluzzii*. A)** Alignment coordinates of *k*-mers that are significantly associated with pirimiphos-methyl resistance in Ivorian *A. coluzzii* samples from the *Anopheles gambiae* 1000 Genomes Phase 2. For each *k*-mer, we report the alignment sequence and coordinates, and whether it overlaps the *Ace1* gene or the *Ace1* duplication. This table includes both primary and secondary alignments as reported by *bwa mem*, as well as non-aligned *k*-mers (chromosome “NA”). **B)** Correlation of alignment frequencies and *Ace1* copy number in each sample, for each of the 409 *k-* mers that mapped to chromosomal arms 2R, 2L, 3R, 3L or X. We report Pearson’s correlation coefficients (*r*) and *p-*values. **C)** Sequences of the 446 assembled *k*-mers that are significantly associated with pirimiphos-methyl resistance. **D)** Frequencies of the 9,603 *k-*mers (before assembly) in each of the Côte d’Ivoire *A. coluzzii* samples (*Anopheles gambiae* 1000 Genomes Phase 2, n=71).

**Supplementary Material SM9. Identification of tagging variants for *Ace1 280S*. A)** Density of phased variants in the *Ace1* duplication region, from *Anopheles gambiae* 1000 Genomes Phase 2. **B)** Density of phased variants within the *Ace1* duplication, focusing on the *Ace1* gene body. **C)** Linkage disequilibrium between the *G280S* alleles and nearby phased variants (Huff and Roger’s *r*), highlighting the coordinates of four nearby variants tightly linked to *280S* alleles, and their frequencies in Côte d’Ivoire *A. coluzzii*. The four linked and phased variants can be used to tag the *280S* haplotype.

**Supplementary Material SM10. Haplotype networks around the *Ace1* tagging variants**. Minimum spanning tree networks built from phased variants located around each of the four tagging variants (panels A to D). Each node in the network is colored according to its population composition (left panels) or linkage to the *280S* or *wt* alleles in *Ace1* (right panels).

**Supplementary Material SM11. Signals of selection in the *Ace1* duplication**. Garud *H* statistics and haplotype diversity in *280S*-linked and *wt-*linked haplotypes in the genomic window around the *Ace1* duplication breakpoints, calculated for each of the main haplotype clusters defined around each of the four tagging variants (panels A to D; haplotype clusters from Supplementary Material SM10). For each tagging variant and duplication breakpoint (upstream/downstream), we report the average value of each statistic and standard errors from sample jack-knifing.

**Supplementary Material SM12. Extended haplotype homozygosity in the *Ace1* duplication**. Extended haplotype homozygosity (*EHH*) of *280S*-linked and *wt-*linked haplotypes in the genomic window around the *Ace1* duplication breakpoints, calculated for each of the main haplotype clusters defined around each of the four tagging variants (panels A to D; haplotype clusters from Supplementary Material SM10). For each tagging variant and duplication breakpoint (upstream/downstream), we report the area under the *EHH* curve (*a*) and the distance around the duplication breakpoint where *EHH*>0.05 and *EHH*>0.95.

**Supplementary Material SM13. Introgression of the *Ace1* duplication**. Patterson *D* statistics to test introgression of the *Ace1* duplication region between various populations of *A. coluzzii* (populations A/B), *A. gambiae* (populations C) and multiple outgroup species (population O). Specifically, we use *A. coluzzii* from Côte d’Ivoire (CIcol), Ghana (GHcol) and Burkina-Faso (BFcol) with and without duplications (labelled as TRUE and FALSE respectively) as populations A; *A. coluzzii* from Angola as population B (always labelled as FALSE); and various *A. gambiae* populations C from Ghana (GHgam), Burkina-Faso (BFgam), Guinea (GNgam), Gabon (GAgam) and Cameroon (CMgam) with and without duplications (TRUE and FALSE labels, respectively); and four different outgroup species (panels A to D: *A. arabiensis, A. melas, A. merus* and *A. quadriannulatus*). For each comparison, we report the average *D* statistic from the *Ace1* duplicated region with standard errors, and *Z-*scores and *p*-values derived from standardised *D* values (unit variance).

**Supplementary Material SM14. Haplotype alignments. A)** Alignment of phased variants from within the *Ace1* duplication region (2R:3,436,800-3,639,600), using 345 samples from West African populations (*Anopheles gambiae* 1000 Genomes) with *Ace1* duplications (Guinea *A. gambiae*, Côte d’Ivoire *A. coluzzii*, Ghana *A. gambiae* and *A. coluzzii*, Burkina Faso *A. gambiae* and *A. coluzzii*). **B)** Id., from a region upstream of the duplication (50 kbp starting at 3,436,800 – 1 Mb). **C)** Id., from a region downstream of the duplication (50 kbp starting at 3,639,600 + 1 Mb).

**Supplementary Material SM15. Haplotype phylogenies**. Maximum-Likelihood phylogenetic analyses of haplotypes from within the *Ace1* duplication (A), upstream (B) and downstream (C) regions. Trees are unrooted. Tips are color-coded according to duplication presence/absence and species. UF bootstrap supports indicated in each node.

