## Supplementary material for "Resistance to pirimiphos-methyl in West African *Anopheles* is spreading via duplication and introgression of the *Ace1* locus": SM1. Homology of Ace1 mutations

**A)** Alignment of *Ace1* orthologs: selected animal sequences

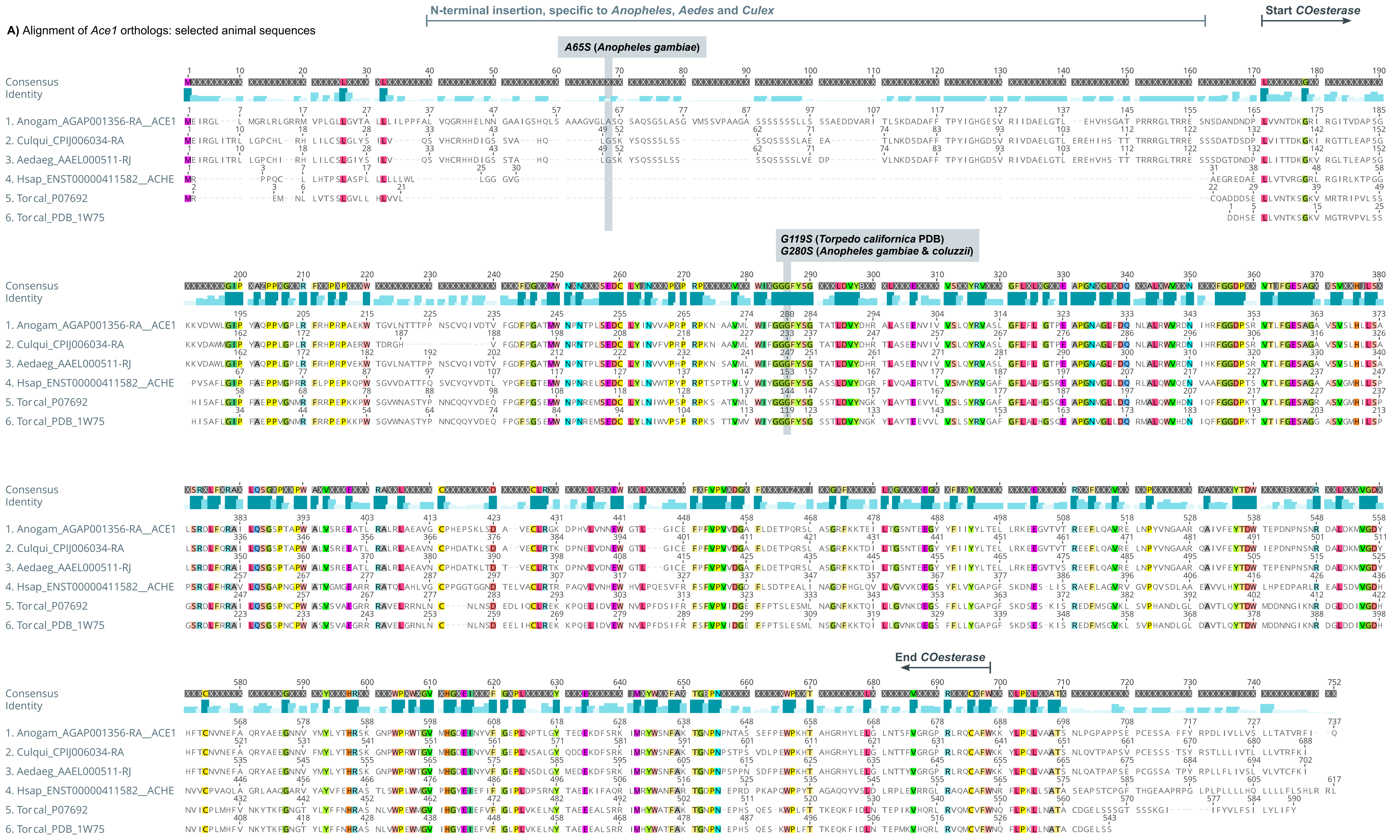


**B)** Alignment of *Ace1* orthologs: *Anopheles*, *Culex* and *Aedes*

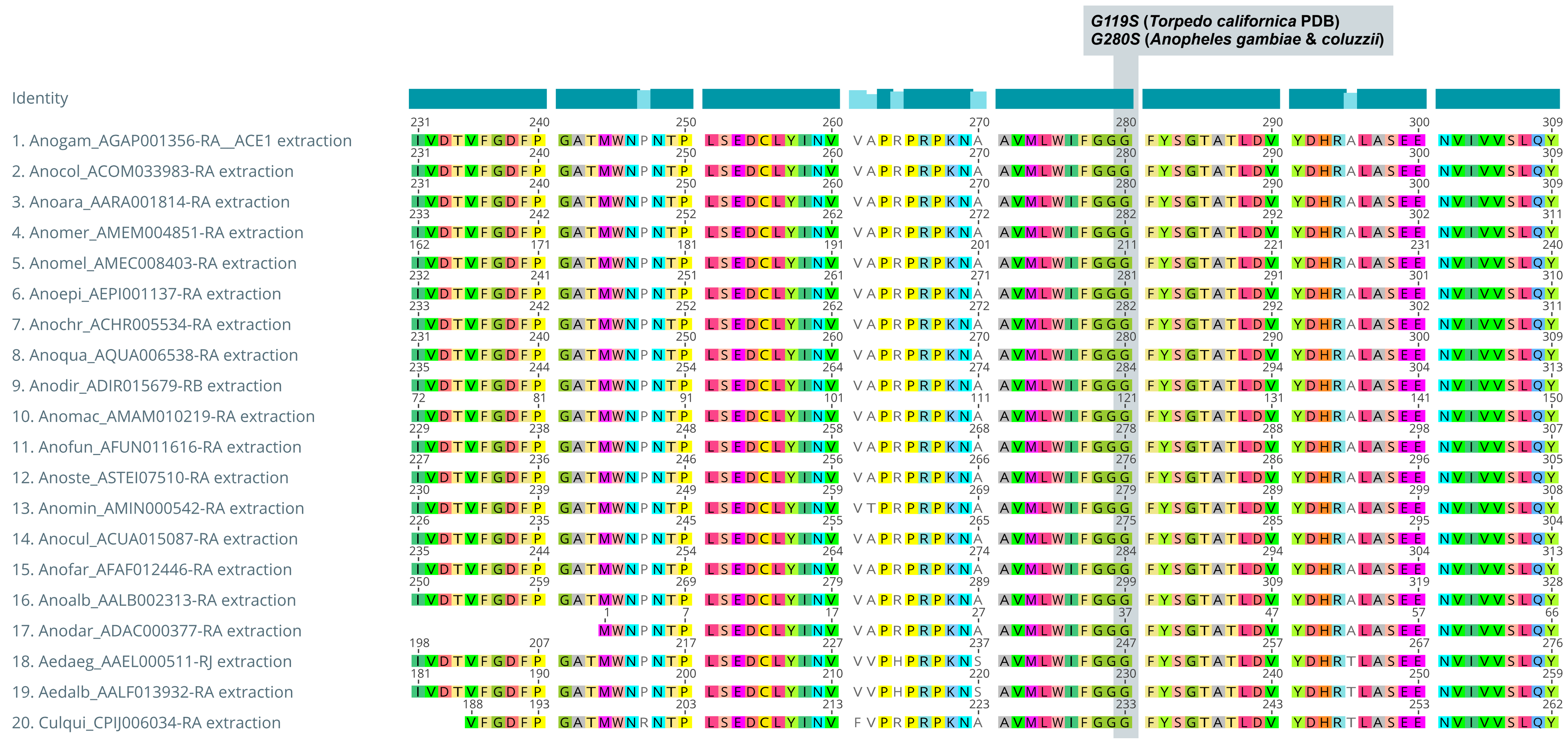


**C)** Phylogenetic analysis Ace homologs

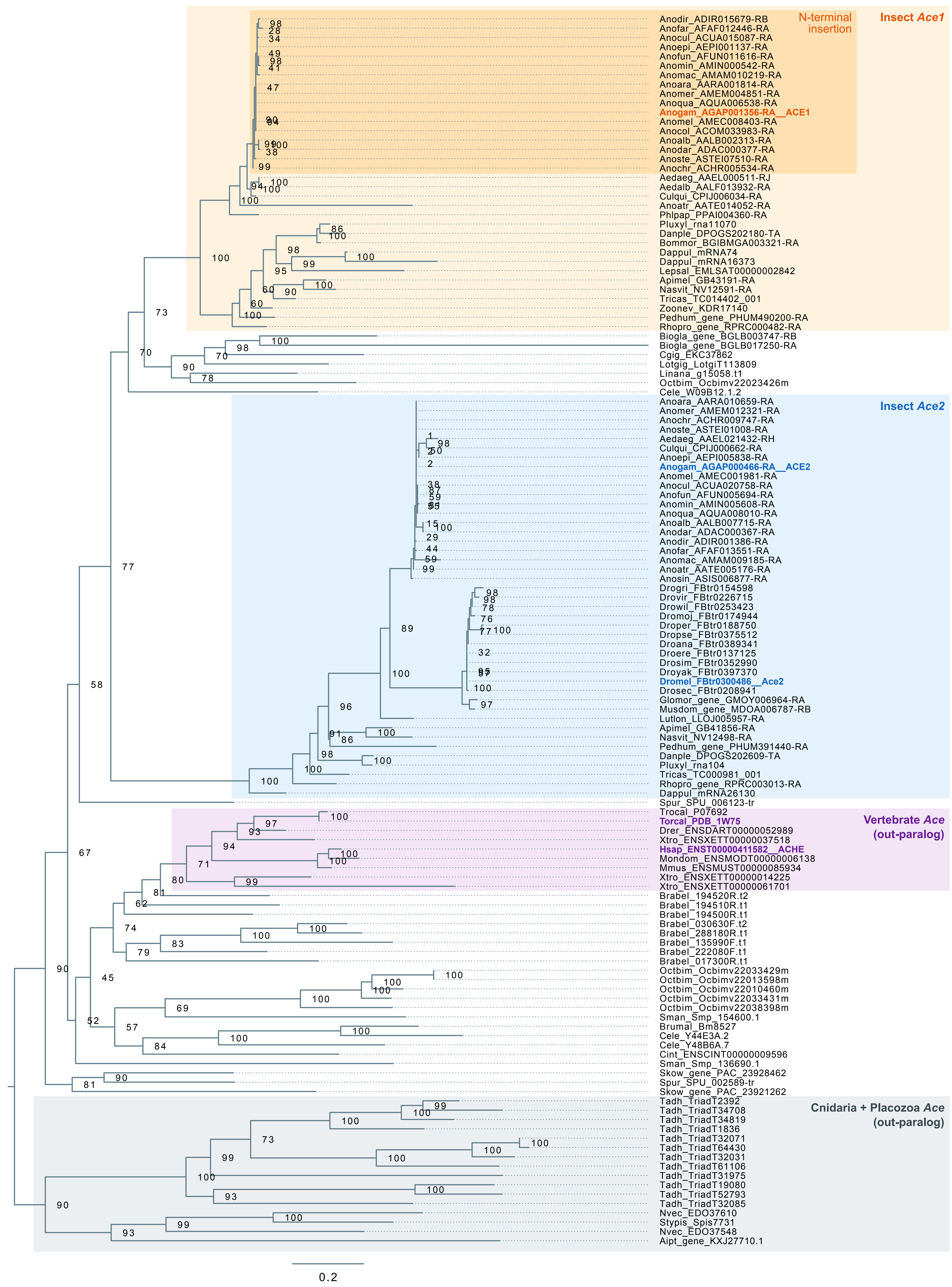
