## Supplementary material for "Resistance to pirimiphos-methyl in West African *Anopheles* is spreading via duplication and introgression of the *Ace1* locus": SM9. Identification of tagging variants for Ace1 280S

**A)** Density of phased variants around *Ace1* duplication region

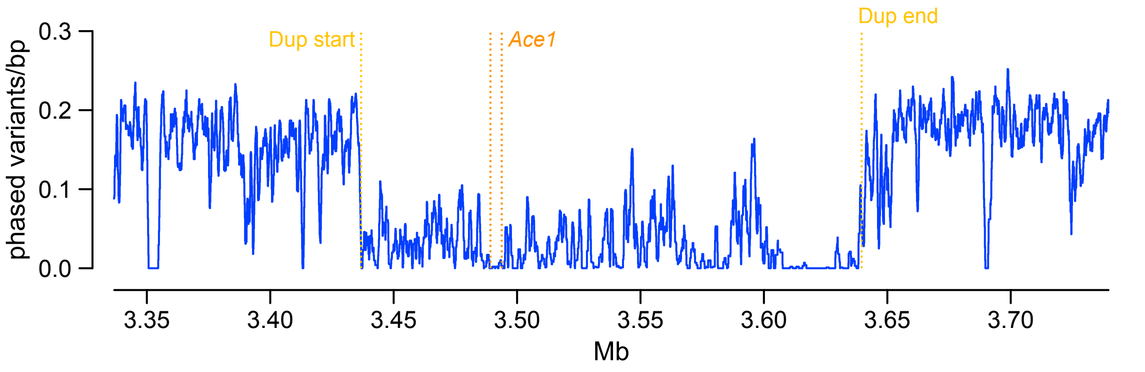


**B)** Density of phased variants within *Ace1* duplication region

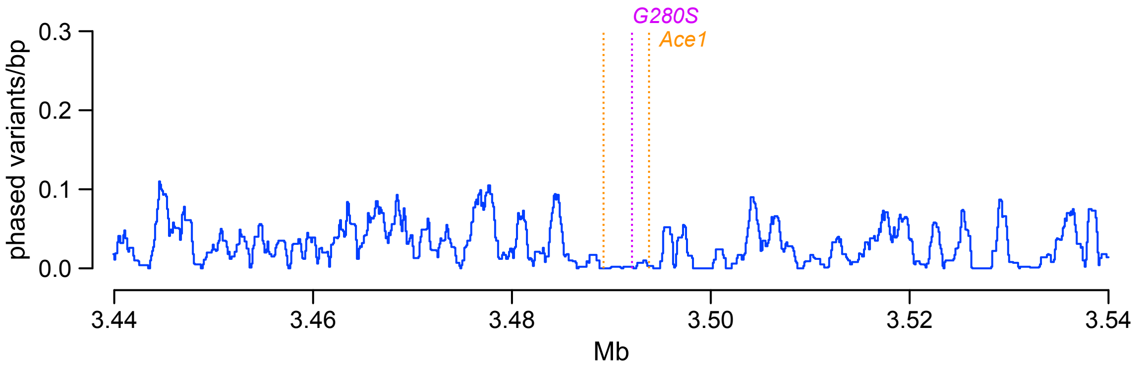


**C)** Linkage disequilibrium between *GG280S* and nearby phased variants

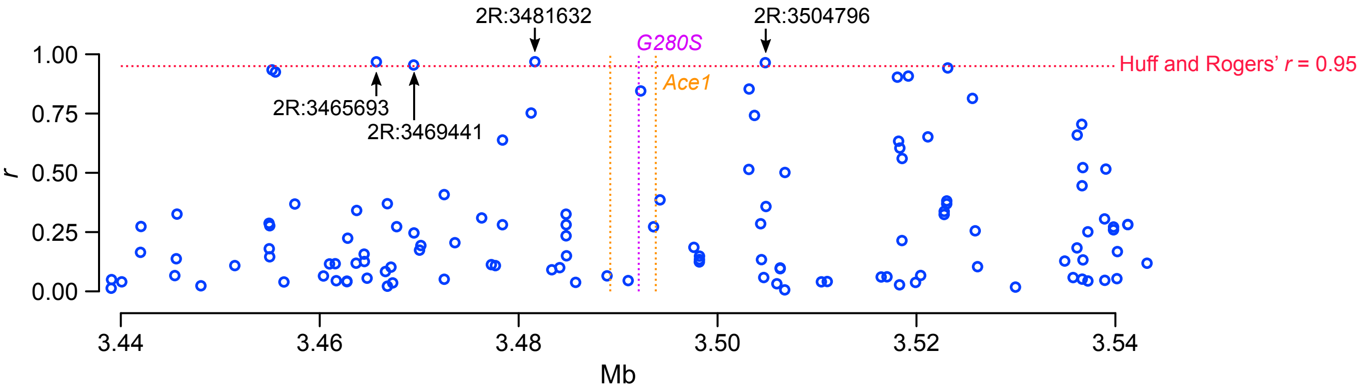


| chrom | pos | LD with G280S | Freq CIcol | Phased? |
|---|---|---|---|---|
| 2R | 3444618 | 0.97264516 | 0.4366 | 0 |
| 2R | 3445910 | 0.9504605 | 0.4366 | 0 |
| 2R | 3447556 | 0.96361303 | 0.4366 | 0 |
| 2R | 3456166 | 0.9675373 | 0.4366 | 0 |
| 2R | 3456937 | 0.97637683 | 0.4366 | 0 |
| 2R | 3459582 | 0.9591772 | 0.4366 | 0 |
| 2R | 3464705 | 0.96810204 | 0.4366 | 0 |
| 2R | 3465693 | 0.96810204 | 0.4366 | 1 *** |
| 2R | 3466268 | 0.9719312 | 0.4366 | 0 |
| 2R | 3469441 | 0.95479333 | 0.4366 | 1 *** |
| 2R | 3469672 | 0.96308875 | 0.4366 | 0 |
| 2R | 3475119 | 0.9658597 | 0.4366 | 0 |
| 2R | 3477809 | 0.98811066 | 0.4366 | 0 |
| 2R | 3478595 | 1.0 | 0.4366 | 0 |
| 2R | 3479474 | 0.98421353 | 0.4366 | 0 |
| 2R | 3480330 | 1.0 | 0.4366 | 0 |
| 2R | 3480406 | 0.98421353 | 0.4366 | 0 |
| 2R | 3481632 | 0.96864766 | 0.4295 | 1 *** |
| 2R | 3482092 | 0.9716037 | 0.4154 | 0 |
| 2R | 3483376 | 0.99236995 | 0.4366 | 0 |
| 2R | 3483539 | 1.0 | 0.4366 | 0 |
| 2R | 3486607 | 0.9959919 | 0.4366 | 0 |
| 2R | 3487859 | 1.0 | 0.4366 | 0 |
| 2R | 3496635 | 0.98388374 | 0.4366 | 0 |
| 2R | 3498622 | 0.96866876 | 0.4366 | 0 |
| 2R | 3498740 | 0.97213244 | 0.4366 | 0 |
| 2R | 3504796 | 0.96453327 | 0.4436 | 1 *** |
| 2R | 3507961 | 0.97605 | 0.4507 | 0 |
