## Supplementary material for "Resistance to pirimiphos-methyl in West African *Anopheles* is spreading via duplication and introgression of the *Ace1* locus": SM10. Haplotype networks around the Ace1 tagging variants

### A) Haplotype networks 2R:3465693

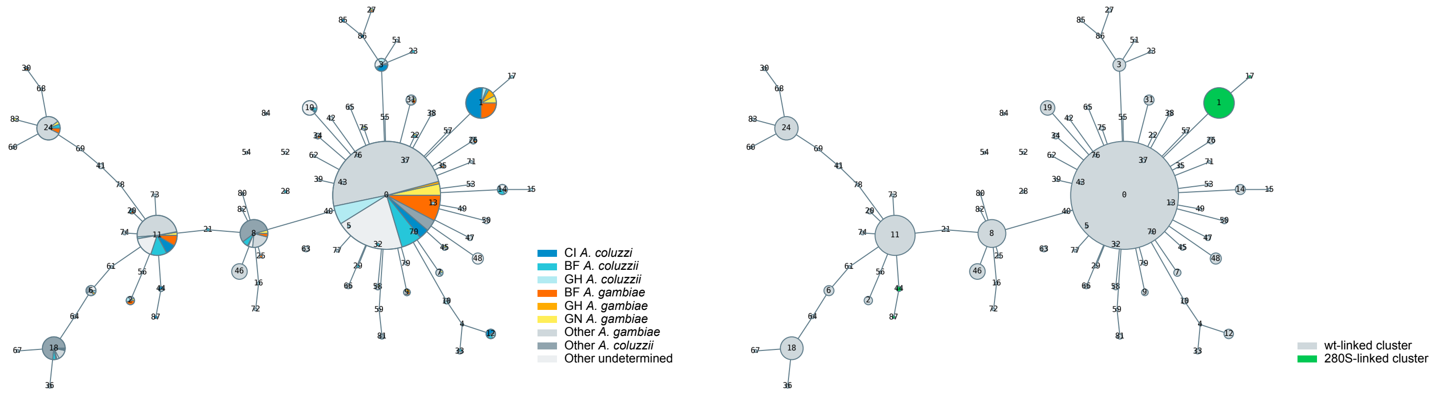


Ace1due 2R:3465693 . AGAP001355-AGA 2R:3465693 (3465693), allele 0
2284 haps clustered into 88 clusters with mst maxdist 1, from 67 phased vars located +/- 300 bp

### B) Haplotype networks 2R:3469441

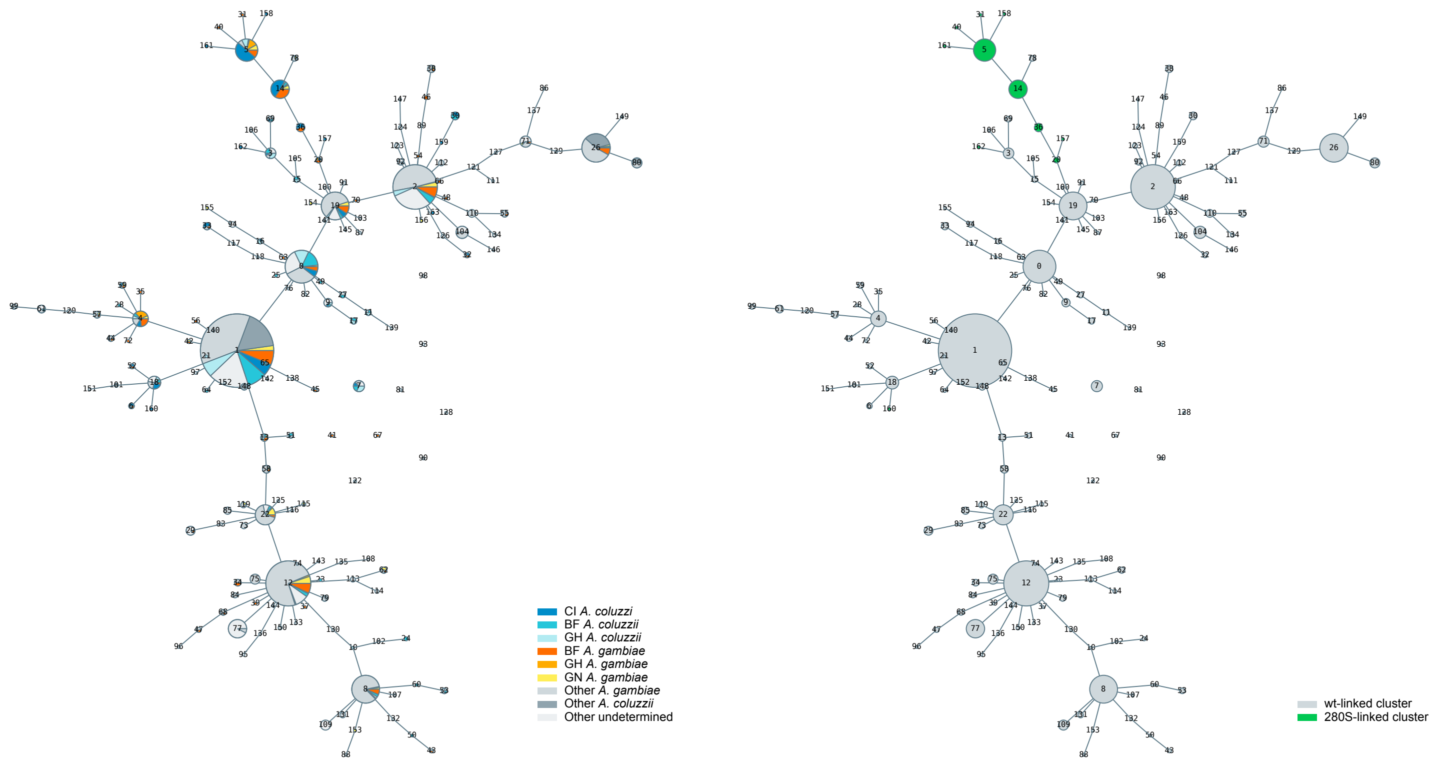


Ace1due 2R:3469441 . AGAP001355-AGA 2R:3469441 (3469441), allele 0
2284 haps clustered into 164 clusters with mst maxdist 1, from 71 phased vars located +/- 300 bp

### C) Haplotype networks 2R:3481632

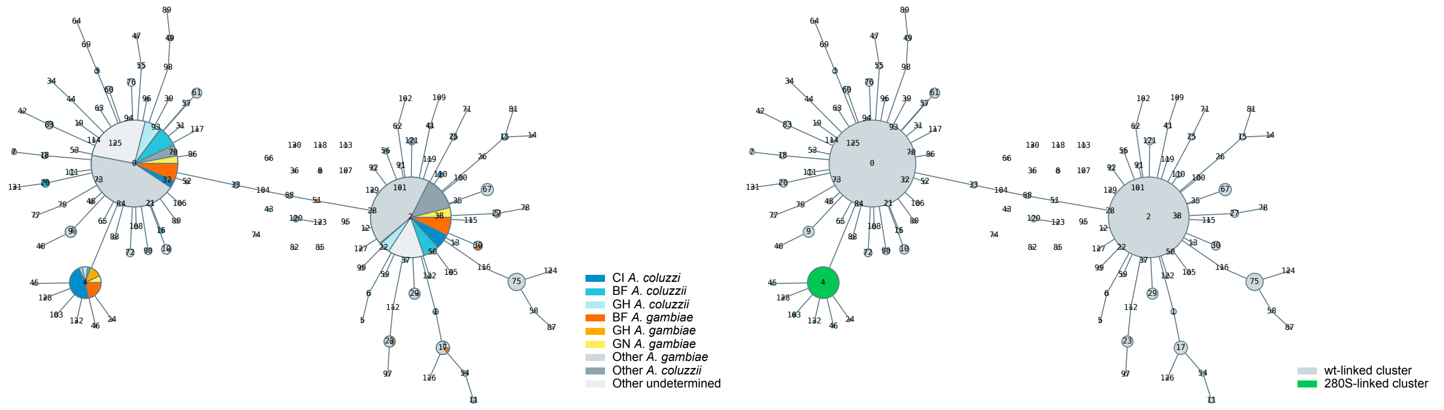


Ace1due 2R:3481632 . AGAP001356 2R:3481632 (3481632), allele 0
2284 haps clustered into 133 clusters with mst maxdist 1, from 104 phased vars located +/- 300 bp

### D) Haplotype networks 2R:3504796

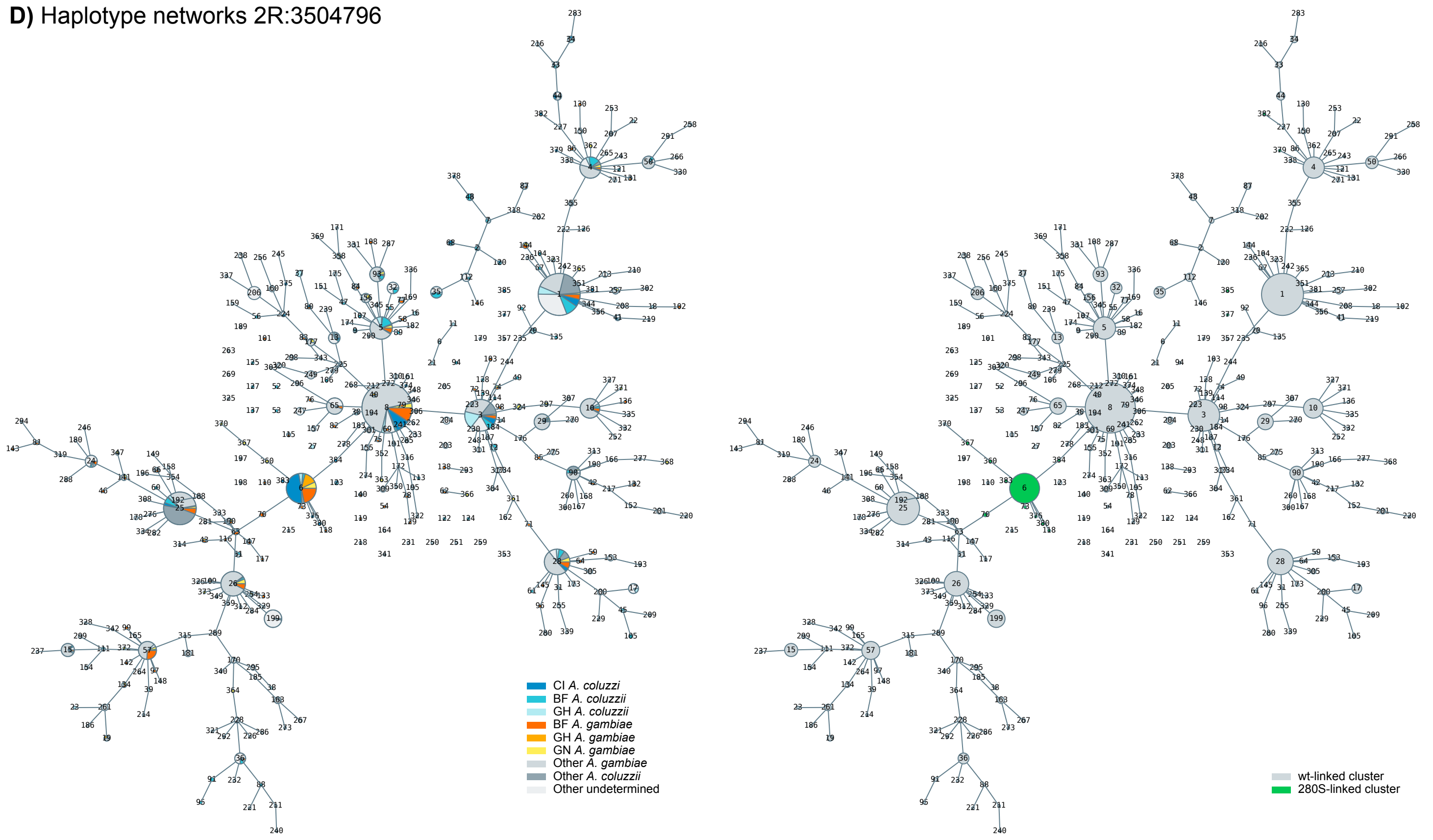


Ace1due 2R:3504796 . AGAP001357 2R:3504796 (3504796), allele 0
2284 haps clustered into 386 clusters with mst maxdist 1, from 164 phased vars located +/- 300 bp
