## Supplementary material for "Resistance to pirimiphos-methyl in West African *Anopheles* is spreading via duplication and introgression of the *Ace1* locus": SM11. Signals of selection in the Ace1 duplication

### **A)** Tagging variant 2R:3465693

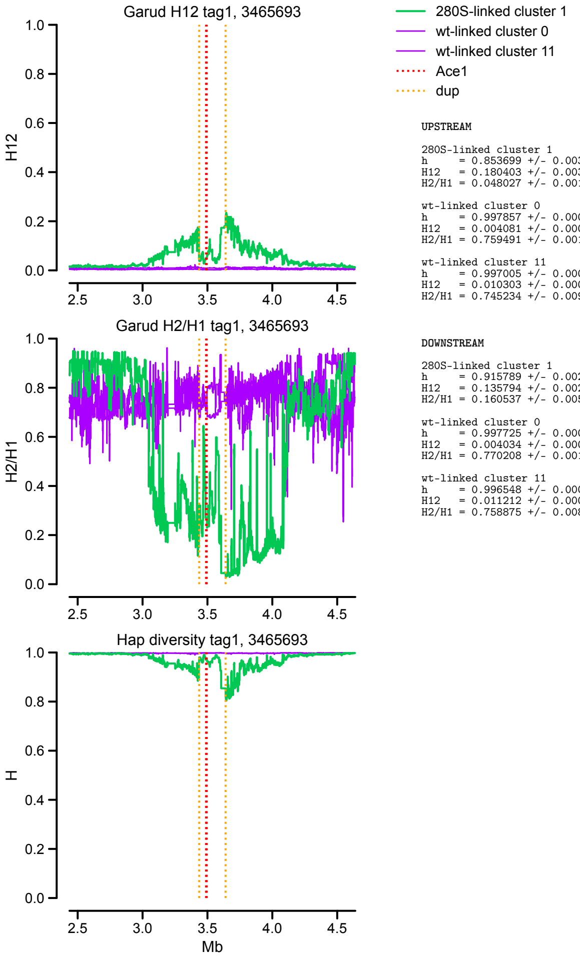


```
UPSTREAM

280S-linked cluster 1
h      = 0.853699 +/- 0.003219
H12    = 0.180403 +/- 0.003551
H2/H1 = 0.048027 +/- 0.001552

wt-linked cluster 0
h      = 0.997857 +/- 0.000008
H12    = 0.004081 +/- 0.000015
H2/H1 = 0.759491 +/- 0.001641

wt-linked cluster 11
h      = 0.997005 +/- 0.000091
H12    = 0.010303 +/- 0.000169
H2/H1 = 0.745234 +/- 0.009099


DOWNSTREAM

280S-linked cluster 1
h      = 0.915789 +/- 0.002090
H12    = 0.135794 +/- 0.002912
H2/H1 = 0.160537 +/- 0.005901

wt-linked cluster 0
h      = 0.997725 +/- 0.000007
H12    = 0.004034 +/- 0.000013
H2/H1 = 0.770208 +/- 0.001570

wt-linked cluster 11
h      = 0.996548 +/- 0.000095
H12    = 0.011212 +/- 0.000187
H2/H1 = 0.758875 +/- 0.008788
```

### **B)** Tagging variant 2R:3469441

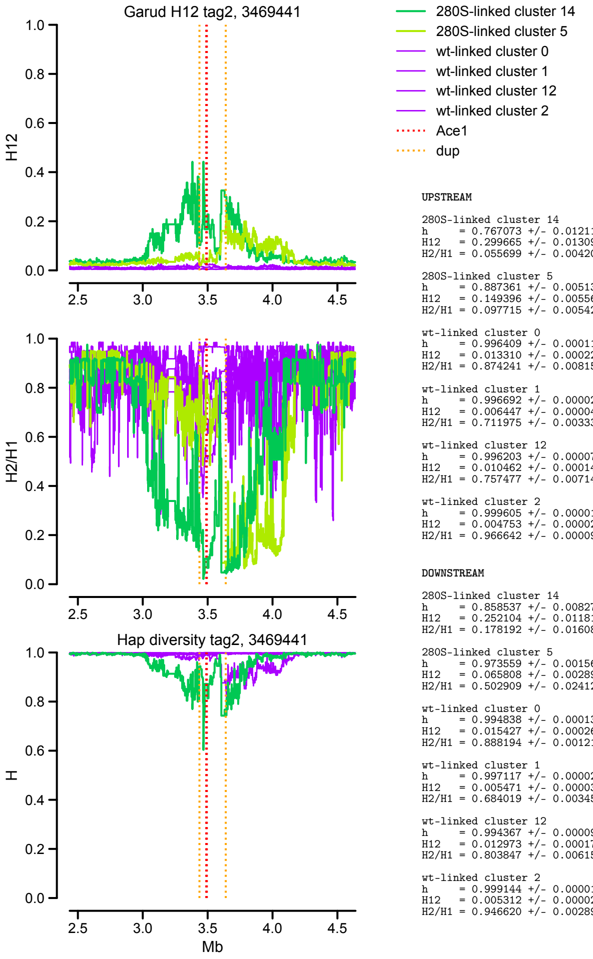


```
UPSTREAM

280S-linked cluster 14
h      = 0.767073 +/- 0.012118
H12    = 0.299665 +/- 0.013090
H2/H1 = 0.055699 +/- 0.004204

280S-linked cluster 5
h      = 0.887361 +/- 0.005137
H12    = 0.149396 +/- 0.005561
H2/H1 = 0.097715 +/- 0.005425

wt-linked cluster 0
h      = 0.996409 +/- 0.000117
H12    = 0.013310 +/- 0.000229
H2/H1 = 0.874241 +/- 0.008154

wt-linked cluster 1
h      = 0.996692 +/- 0.000024
H12    = 0.006447 +/- 0.000042
H2/H1 = 0.711975 +/- 0.003330

wt-linked cluster 12
h      = 0.996203 +/- 0.000075
H12    = 0.010462 +/- 0.000148
H2/H1 = 0.757477 +/- 0.007142

wt-linked cluster 2
h      = 0.999605 +/- 0.000012
H12    = 0.004753 +/- 0.000026
H2/H1 = 0.966642 +/- 0.000092


DOWNSTREAM

280S-linked cluster 14
h      = 0.858537 +/- 0.008278
H12    = 0.252104 +/- 0.011815
H2/H1 = 0.178192 +/- 0.016087

280S-linked cluster 5
h      = 0.973559 +/- 0.001565
H12    = 0.065808 +/- 0.002894
H2/H1 = 0.502909 +/- 0.024124

wt-linked cluster 0
h      = 0.994838 +/- 0.000136
H12    = 0.015427 +/- 0.000268
H2/H1 = 0.888194 +/- 0.001215

wt-linked cluster 1
h      = 0.997117 +/- 0.000021
H12    = 0.005471 +/- 0.000034
H2/H1 = 0.684019 +/- 0.003451

wt-linked cluster 12
h      = 0.994367 +/- 0.000090
H12    = 0.012973 +/- 0.000173
H2/H1 = 0.803847 +/- 0.006152

wt-linked cluster 2
h      = 0.999144 +/- 0.000019
H12    = 0.005312 +/- 0.000028
H2/H1 = 0.946620 +/- 0.002898
```

### **C)** Tagging variant 2R:3481632

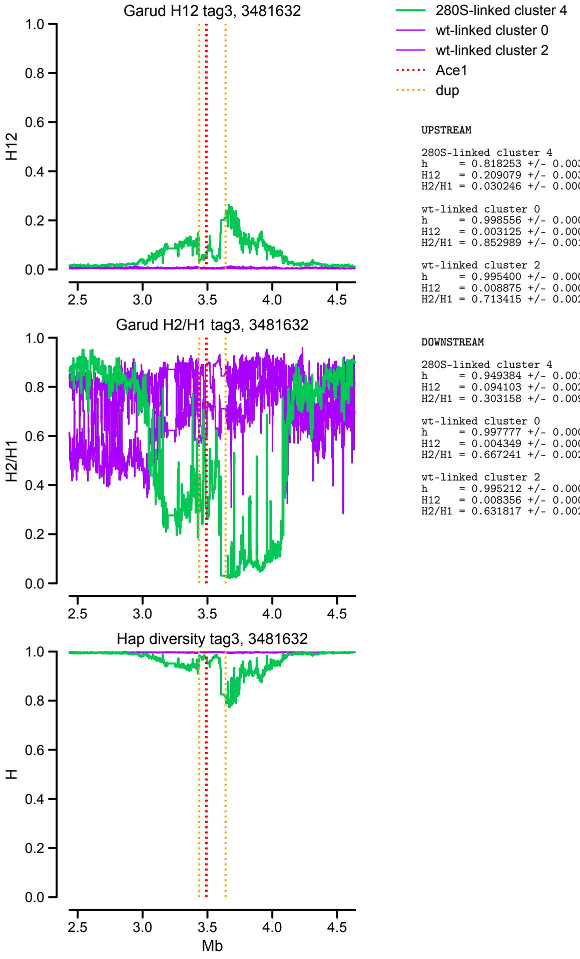


```
UPSTREAM

280S-linked cluster 4
h      = 0.818253 +/- 0.003414
H12    = 0.209079 +/- 0.003609
H2/H1 = 0.030246 +/- 0.000840

wt-linked cluster 0
h      = 0.998556 +/- 0.000008
H12    = 0.003125 +/- 0.000014
H2/H1 = 0.852989 +/- 0.001952

wt-linked cluster 2
h      = 0.995400 +/- 0.000026
H12    = 0.008875 +/- 0.000051
H2/H1 = 0.713415 +/- 0.002716


DOWNSTREAM

280S-linked cluster 4
h      = 0.949384 +/- 0.001251
H12    = 0.094103 +/- 0.002064
H2/H1 = 0.303158 +/- 0.009604

wt-linked cluster 0
h      = 0.997777 +/- 0.000013
H12    = 0.004349 +/- 0.000022
H2/H1 = 0.667241 +/- 0.002710

wt-linked cluster 2
h      = 0.995212 +/- 0.000026
H12    = 0.008356 +/- 0.000045
H2/H1 = 0.631817 +/- 0.002834
```

### **D)** Tagging variant 2R:3504796

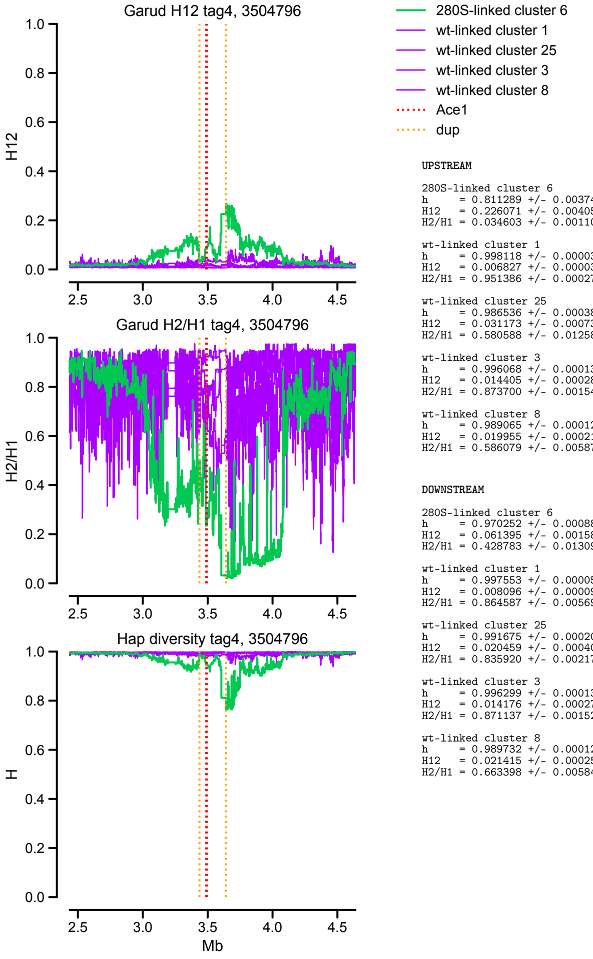


```
UPSTREAM

280S-linked cluster 6
h      = 0.811289 +/- 0.003744
H12    = 0.226071 +/- 0.004056
H2/H1 = 0.034603 +/- 0.001104

wt-linked cluster 1
h      = 0.998118 +/- 0.000034
H12    = 0.006827 +/- 0.000034
H2/H1 = 0.951386 +/- 0.000270

wt-linked cluster 25
h      = 0.986536 +/- 0.000385
H12    = 0.031173 +/- 0.000730
H2/H1 = 0.580588 +/- 0.012581

wt-linked cluster 3
h      = 0.996068 +/- 0.000138
H12    = 0.014405 +/- 0.000280
H2/H1 = 0.873700 +/- 0.001543

wt-linked cluster 8
h      = 0.989065 +/- 0.000126
H12    = 0.019955 +/- 0.000217
H2/H1 = 0.586079 +/- 0.005876


DOWNSTREAM

280S-linked cluster 6
h      = 0.970252 +/- 0.000886
H12    = 0.061395 +/- 0.001583
H2/H1 = 0.428783 +/- 0.013098

wt-linked cluster 1
h      = 0.997553 +/- 0.000050
H12    = 0.008096 +/- 0.000094
H2/H1 = 0.864587 +/- 0.005691

wt-linked cluster 25
h      = 0.991675 +/- 0.000202
H12    = 0.020459 +/- 0.000400
H2/H1 = 0.835920 +/- 0.002176

wt-linked cluster 3
h      = 0.996299 +/- 0.000131
H12    = 0.014176 +/- 0.000277
H2/H1 = 0.871137 +/- 0.001528

wt-linked cluster 8
h      = 0.989732 +/- 0.000122
H12    = 0.021415 +/- 0.000250
H2/H1 = 0.663398 +/- 0.005848
```
