## Supplementary material for "Resistance to pirimiphos-methyl in West African *Anopheles* is spreading via duplication and introgression of the *Ace1* locus": SM12. Extended haplotype homozygosity in the Ace1 duplication

### **A)** Tagging variant 2R:3465693

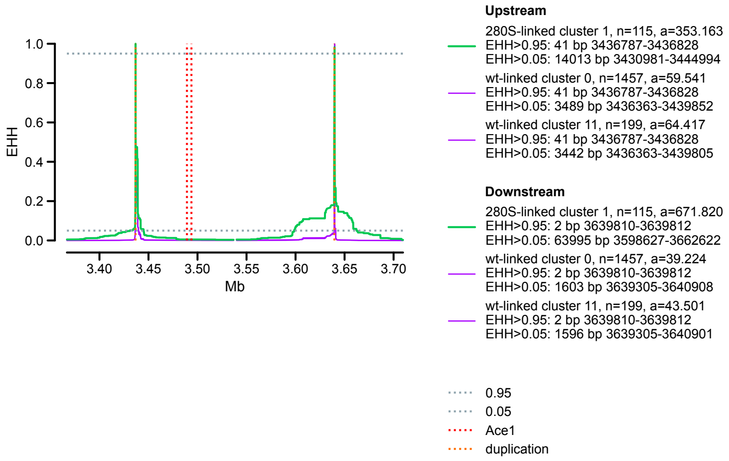


### **B)** Tagging variant 2R:3469441

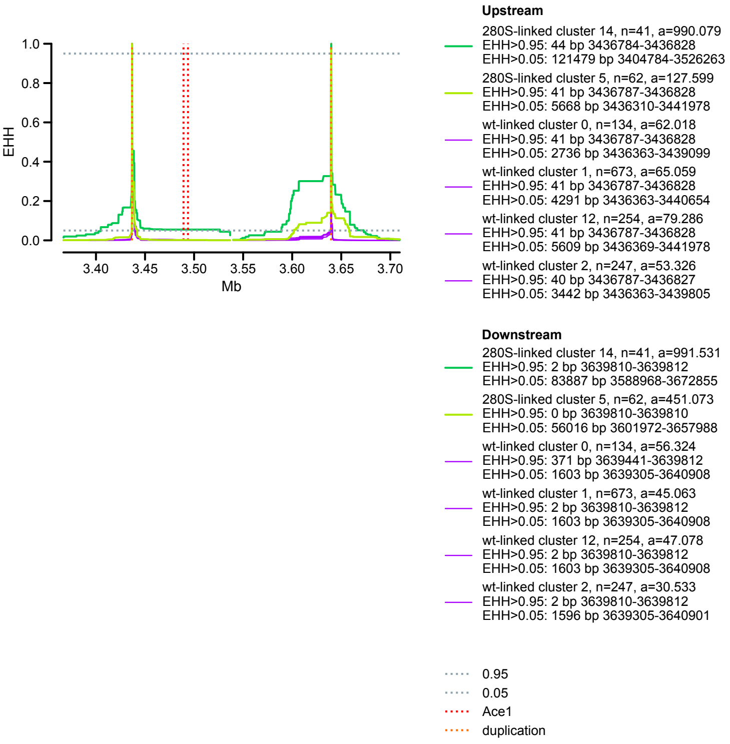


### **C)** Tagging variant 2R:3481632

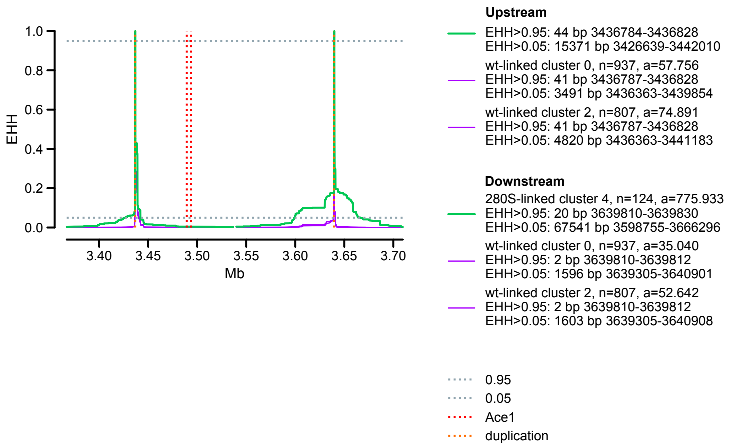


### **D)** Tagging variant 2R:3504796

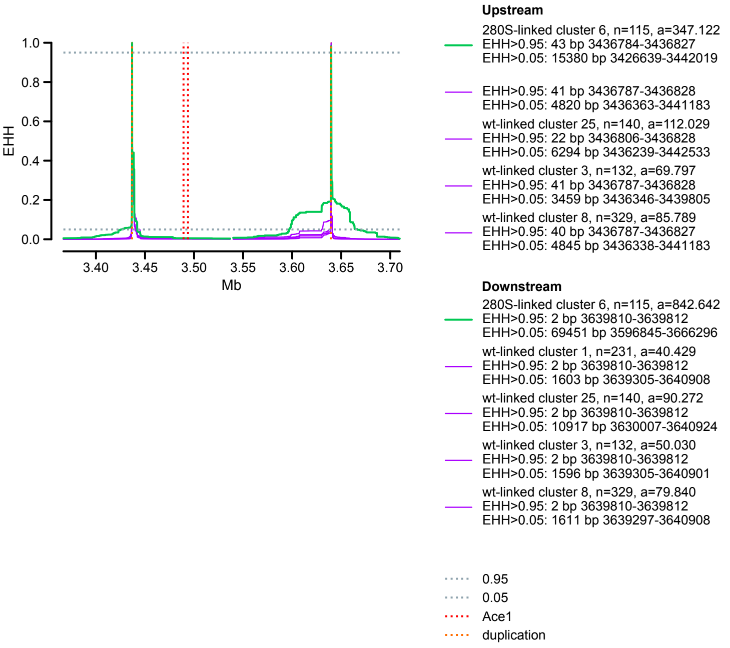
