## Supplementary material for "Resistance to pirimiphos-methyl in West African *Anopheles* is spreading via duplication and introgression of the *Ace1* locus": SM13. Introgression of the Ace1 duplication

### **A)** Introgression in duplicated region, D=*A. arabiensis*

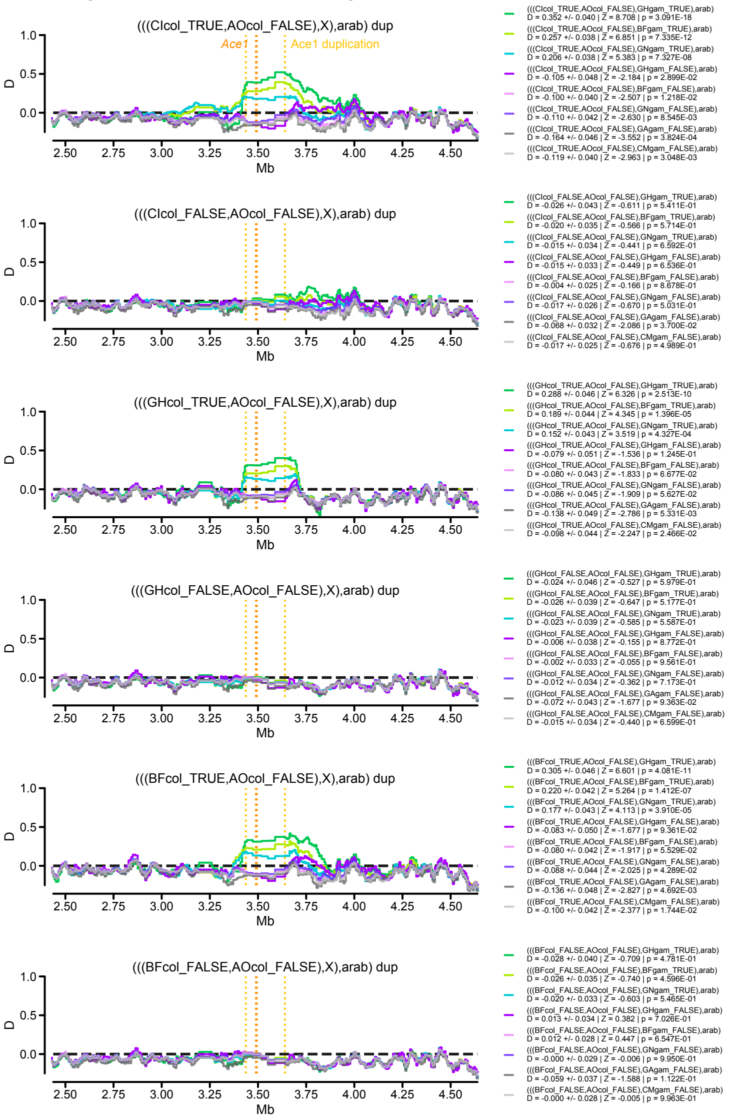


### **B)** Introgression in duplicated region, D=*A. melas*

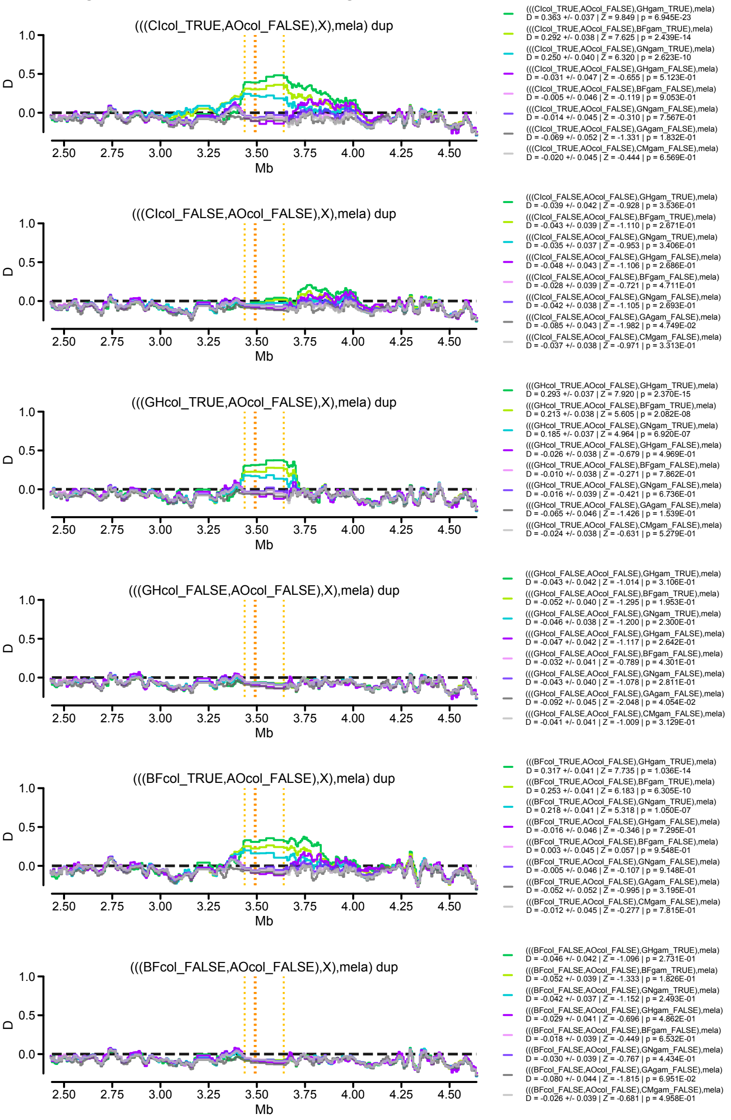


### **C)** Introgression in duplicated region, D=*A. merus*

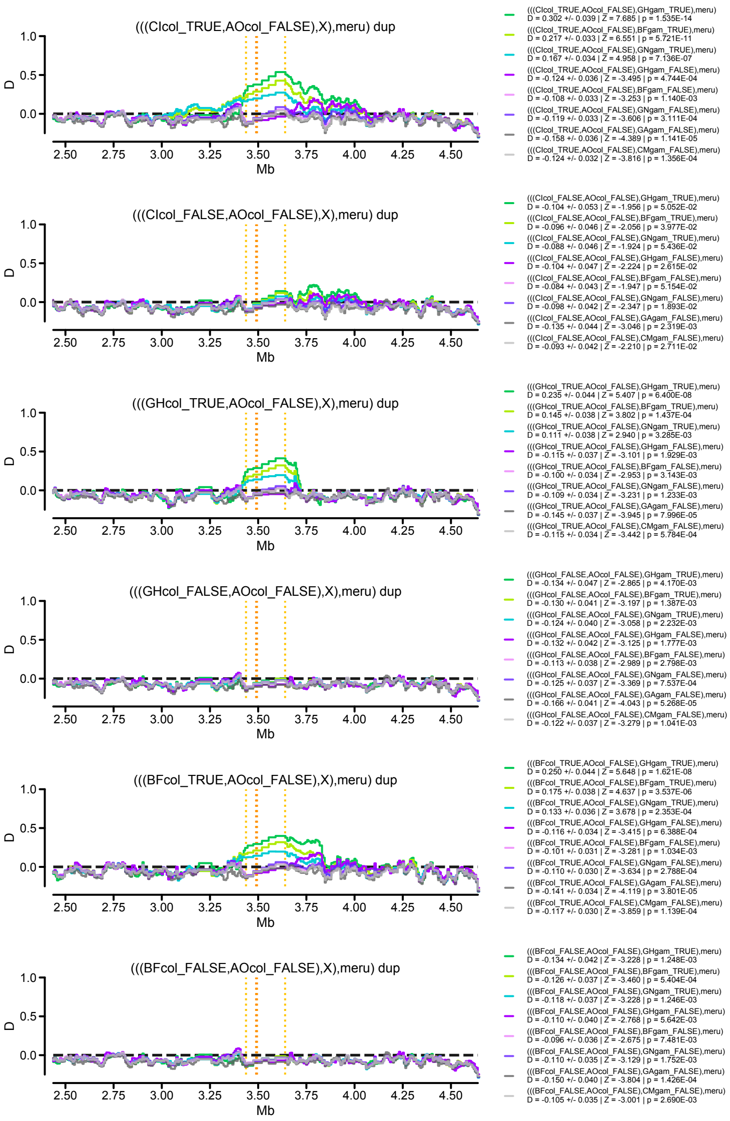


### **D)** Introgression in duplicated region, D=*A. quadriannulatus*

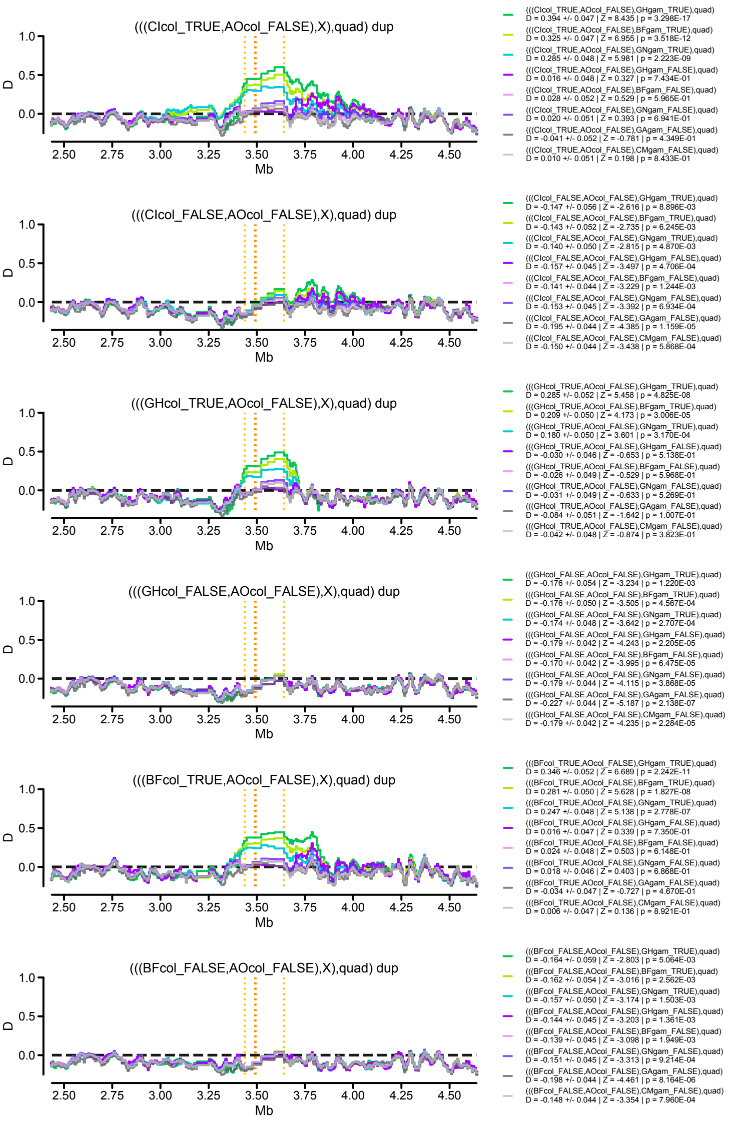
