## Supplementary material for "Resistance to pirimiphos-methyl in West African *Anopheles* is spreading via duplication and introgression of the *Ace1* locus": SM15. Haplotype phylogenies



**A)** Duplication

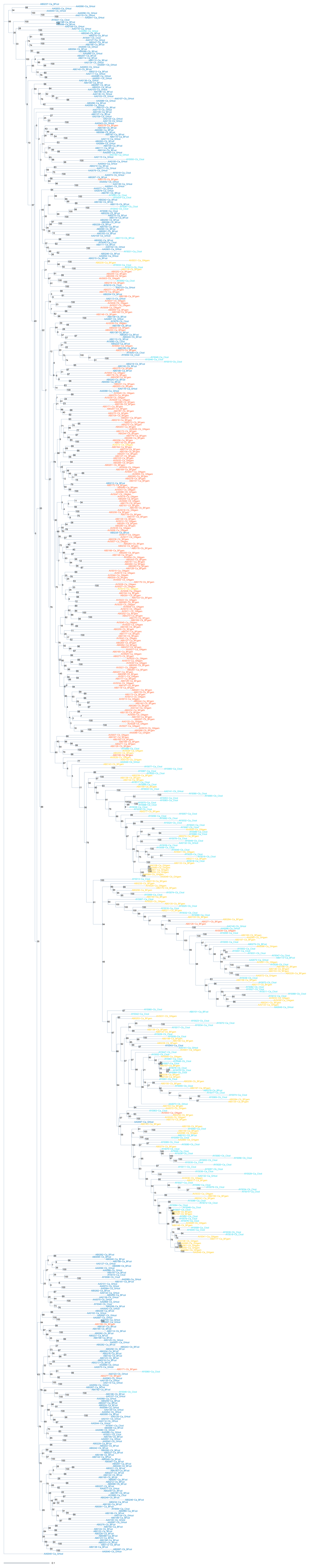

### Supplementary Material SM15

**B)** Upstream

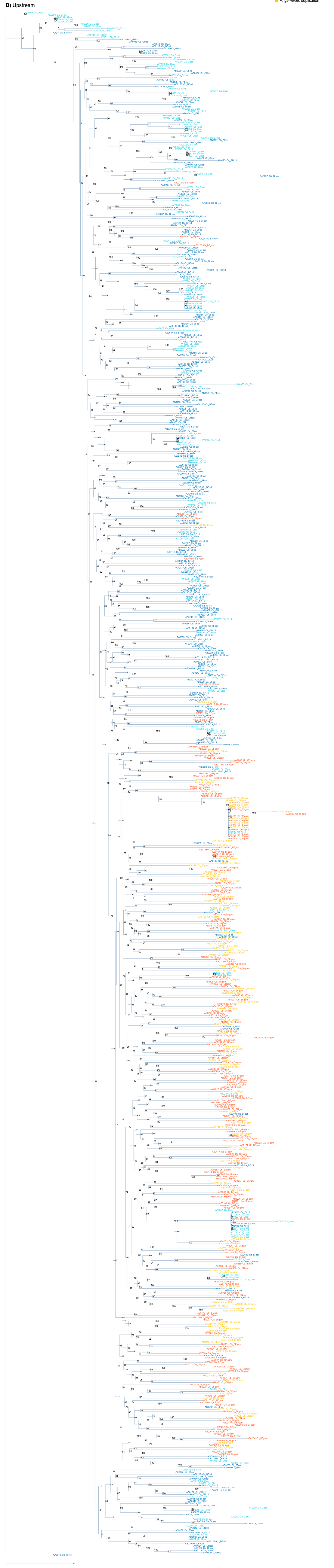

**Supplementary Material SM15**

**C)** Downstream

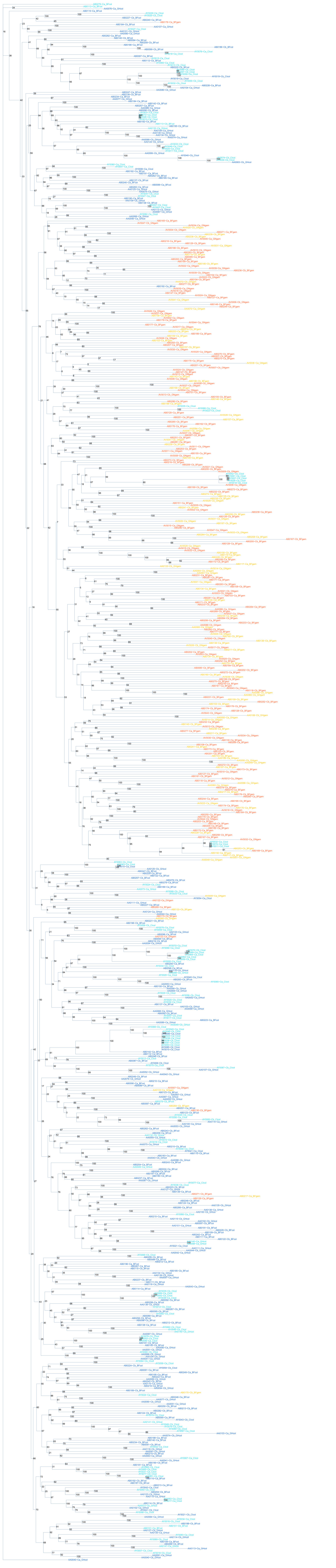
